# From sporulation to village differentiation: the shaping of the social microbiome over rural-to-urban lifestyle transition in Indonesia

**DOI:** 10.1101/2025.02.12.637485

**Authors:** Clarissa A. Febinia, Hirzi Luqman, Pradiptajati Kusuma, Lidwina Priliani, Joseph Lewis, Desak Made Wihandani, Gde Ngurah Indraguna Pinatih, Herawati Sudoyo, Alexandre Almeida, Safarina G. Malik, Guy S. Jacobs

**Author notes:** These authors contributed equally (Clarissa A. Febinia, Hirzi Luqman).

## Abstract

Despite established roles in human health and profound global diversity, existing gut microbiome datasets are biased toward Western urban cohorts, with especial under-representation of Southeast Asia. Here, we present a novel gut microbiome dataset from 116 Indonesians representing a cline from transitional hunter-gatherers to rural agricultural to urban lifestyles. We identify 1,304 species and 3,258 subspecies by assembling 11,070 metagenome-assembled genomes, revealing substantial species (15%) and subspecies level (50%) novelty. Novel taxa are rare, often village-specific, and depleted for sporulation genes, revealing a direct link between bacterial physiology, transmission, prevalence and discovery. We identify a rural-to-urban cline across multiple levels of biological organisation, from species abundance to microbiome composition and diversity. Furthermore, between-community, but not within-community, variation in diet is strongly predictive of microbiome composition, offering compelling evidence that microbiome divergence is driven by community-level differences. Our work highlights the interplay of host lifestyle, population structure and bacterial physiology in shaping microbiome diversity and biogeography, at the key scale of human communities.

## Introduction

The gut microbiome is increasingly recognised as playing important roles in human health and disease (Sonnenburg & Sonnenburg, 2019; Valdes et al., 2018). Yet, despite over a decade of sequencing effort, knowledge of extant diversity is limited as demonstrated by the vast numbers of novel species that continue to be discovered (Carter et al., 2023; Leviatan et al., 2022). Further, research is strongly focussed on urban, industrialised samples from a handful of over-represented countries (Abdill et al., 2022) known to harbour limited microbiome variation with depletion of various phylogenetic groups (Carter et al., 2023; Rosas-Plaza et al., 2022; Yatsunenko et al., 2012). As such, descriptions of the human microbiome, while increasingly extensive (Almeida et al., 2021; Gounot et al., 2022; Pasolli et al., 2019), remain limited and exhibit significant regional and phylogenetic bias. The need to diversify sampling is clear and immediate (Abdill et al., 2022).

Within this context, the Southeast Asian archipelago country of Indonesia has the fourth largest population globally at over 270 million people and is among the most diverse regions worldwide, with over a thousand ethnic groups officially recognised across the archipelago (Na’im & Syaputra, 2012). Despite this, Indonesia is represented by only 0.01% of global metagenomic samples (Abdill et al., 2022). About half of the population live in rural villages, though the country is experiencing rapid growth and urbanisation. The breadth of lifestyles represented, as well as the pace of change, represent clear opportunities to study the nature of lifestyle-associated microbiome transitions, and to broaden representation in microbial datasets.

The impact of lifestyle transitions on the microbiome has been studied in a range of global contexts. Broadly, urban populations show profoundly reduced microbiome diversity and divergent microbial composition, e.g. an excess of *Bacteroides* relative to *Prevotella* and *Treponema*, compared to transitional and rural communities (Gomez et al., 2016; Obregon-Tito et al., 2015; Schnorr et al., 2014). Certain hunter-gatherer groups are particularly diverse (Carter et al., 2023), potentially reflecting microbiome states less affected by modern (agricultural and industrial associated) perturbations and potentially more in line with our evolutionary past. Detailed work on migrant communities (Vangay et al., 2018) has been especially productive in revealing the timescales of microbiome transitions. However, transition is not a single axis, and associations between subsistence strategy and microbiome composition depend on various lifestyle factors and can vary between populations and studies (Gupta et al., 2017; Rosas-Plaza et al., 2022), interacting with numerous drivers to shape microbiome variation in complex ways.

Transitions ultimately arise from the behaviour of individuals living within communities, in the context of microbial diversity that may also be geographically structured (Andreu-Sánchez et al., 2025; Suzuki et al., 2022). At the individual level, environment is a key driver of microbiome composition (Rothschild et al., 2018), along with diet (Wilson et al., 2020; Zhernakova et al., 2016), antibiotic use (Korpela et al., 2016; Nagata et al., 2022; Zhernakova et al., 2016), birth practices (Penders et al., 2006), and age (Badal et al., 2020; Rampelli et al., 2023; Yatsunenko et al., 2012). Communities are increasingly a focus of research, with social interactions and transmission important drivers of strain diversity and sharing (Brito et al., 2019; Valles-Colomer et al., 2023) together shaping the ‘social microbiome’ (Debray et al., 2024), defined as the collective microbial metacommunity of a social group or network (Sarkar et al., 2020, 2024). Yet the relationship between individual behaviour and community living – and indeed, how this interacts with the diversity of microbial physiologies and transmission routes – is hard to disentangle and only partially understood.

In this study, we build a region-specific gut microbiome database by sequencing faecal samples from 116 individuals from 8 communities (Figure 1A, Supplementary Table S1). These communities occupy the islands of Bali, Borneo and New Guinea, spanning an archipelago of approximately 2,500 km yet united by a tropical climate and interconnected recent and more ancient human histories. Critically, the groups also span a range of lifestyles, including transitional hunter-gatherers (Punan Batu), fisher-horticulturalists (Asmat Daikot), mixed forager-agriculturalists (Punan Aput and Punan Tubu Hulu), horticulturalists (Basap), rural agriculturalists (Balinese in Pedawa), peri-urban residents (Punan Tubu Respen) and fully urban residents (Balinese in Denpasar). The Punan (or Penan) traditionally practised a hunter-gatherer lifestyle; the Punan groups in this study hence include communities at different stages of integration with the market economy. The combination of geographic and lifestyle sampling with village-scale microbiome sampling, coupled with genome-resolved metagenomic methods, enables us to identify and explore key drivers of local microbiome diversity. Finally, we describe the social biogeography of microbiome variation and novelty, identifying key bacterial traits associated with microbe geographic distribution and prevalence, with key implications for the understanding of global microbiome diversity.

**Figure 1.**
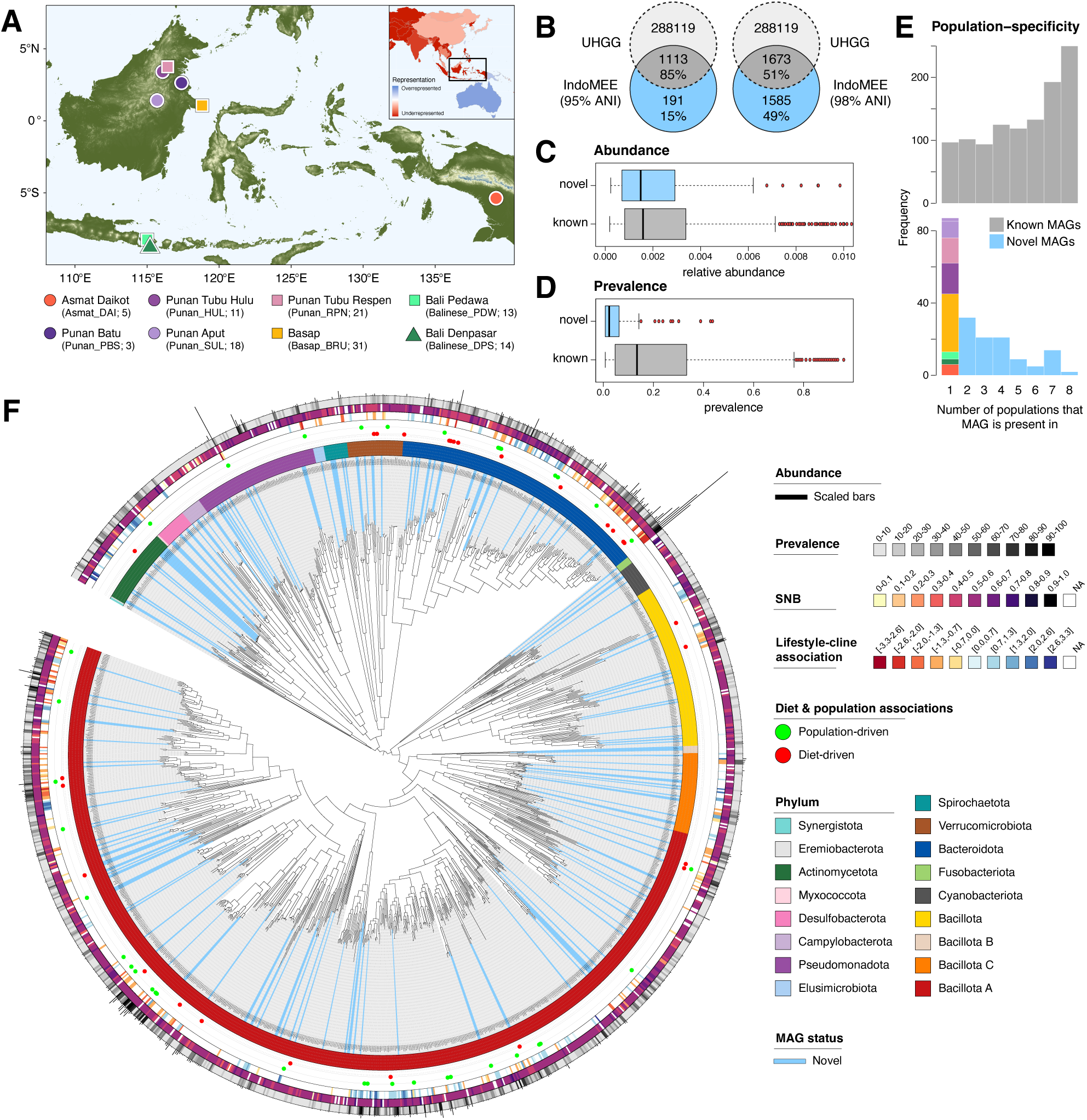
Novel Indonesian microbiome database captures substantial microbial diversity and novelty. **A**) Map of sampled Indonesian communities, with shapes denoting broad lifestyle categories (circles - remote; squares - rural; triangle - urban), and population abbreviations and sample sizes shown in parentheses. Inset map shows countries’ representation in current human microbiome datasets, relative to its share of the world population. Red and blue colours denote countries that are over-represented and under-represented relative to their population size, respectively. Countries with zero samples are indicated in dark red (modified from Abdill, Adamowicz and Blekhman, 2022). **B**) Numbers of novel (blue) and previously described (i.e., known; dark gray) MAGs in the newly constructed IndoMEE gut microbiome reference database, in comparison to the global Unified Human Gastrointestinal Genome (UHGG) database. **C**) Mean within-individual relative abundance, calculated across samples in which MAG is present, of novel MAGs versus known MAGs, shown for species-level MAGs (i.e. 95% ANI). No significant difference is observed between the two categories (Mann-Whitney two-sided test, *p =* 0.15). **D**) Prevalence of novel MAGs compared to known MAGs, shown for species-level MAGs. Novel MAGs are significantly less prevalent (Mann-Whitney two-sided test, *p* < 0.0001) than known MAGs. In box plots (**C** and **D**), centre lines denote medians, box edges indicate lower and upper quartiles, and whiskers extend to 1.5 times the interquartile range. **E**) Histograms of village-level prevalence for novel (bottom; coloured) and known (top; gray) species-level MAGs. Novel MAGs are, relatively, more often unique to a single village (population) than known MAGs in the IndoMEE database (two-sample Kolmogorov-Smirnov test, D = 0.47, p < 0.0001). Colours for village-specific novel MAGs correspond to that in panel A. **F**) Phylogenetic tree of IndoMEE MAGs annotated with status (novel or known; blue and gray radial lines respectively), and (from inner to outer rings) i) phylum, ii) whether they are significantly associated with diet or population, iii) association strength with remote-rural-urban lifestyle cline (if significant), iv) social niche breadth (SNB), v) prevalence, and vi) abundance (Supplementary Table S3).

## Results

### Novel Indonesian microbiome data significantly expands the number of genomes described in Southeast Asia

To characterise regional microbiome diversity, we first constructed a gut microbiome reference database based on faecal samples from our sample of 116 individuals, that were 250 bp paired-end sequenced (average yield = 10.63 Gb, Supplementary Table S2). We combined individual- and co-assembly methods to construct an assembly that was both sensitive to rare taxa and robust against assembly artefacts due to fine-scale genomic variation (Delgado & Andersson, 2022). This yielded 8,633 and 2,437 metagenome-assembled genomes (MAGs) respectively, and 11,070 MAGs total, hereafter referred to as the Indonesian Microbiome Ecology and Evolution (IndoMEE) reference database. This significantly expands the number of microbial genomes described in Southeast Asia (313 and 4497 Southeast Asian MAGs previously identified in Almeida *et al*. (Almeida et al., 2021) and Gounot *et al*. (Gounot et al., 2022), respectively). We combined and dereplicated the individual- and co-assemblies to generate the final reference assembly, which comprised 1,304 (1,296 bacteria, 8 archaea) and 3,258 (3,247 bacteria, 11 archaea) species and subspecies level MAGs, respectively. To classify MAGs, we used GDTB-tk to assign taxonomies based on the Genome Taxonomy Database (GTDBtk release 214). We identify a diverse set of bacterial species in IndoMEE, spread across 17 phyla, 23 classes, 50 orders, 111 families, and 450 genera. Bacillota A is, by far, the most numerous phylum represented (∼50% MAGs), followed by Bacteroidota, Bacillota and Pseudomonadota.

### IndoMEE effectively captures regional diversity, and harbours many village-specific novel microbial species

To assess how well IndoMEE captures regional diversity, we evaluated and compared its mapping rate against state-of-the-art global and regional gut microbiome reference genome databases, namely the Unified Human Gastrointestinal Genome (UHGG) v2.0.1 (Almeida et al., 2021) and Singapore Platinum Metagenomes Project (SPMP) databases (Gounot et al., 2022). Despite being assembled from a similar sample size (n = 116 for IndoMEE vs 109 for SPMP), IndoMEE significantly outperformed SPMP when applied on our Indonesian samples, mapping on average 72.7% of reads compared to 57.4% for the latter (Supplementary Figure S1A). This highlights both the region-specificity of regional databases, and the higher diversity of microbiota captured in IndoMEE (based on diverse Indonesian villages; 1,304 species-level MAGs) compared to SPMP (urban Singapore; 685 species-level MAGs). IndoMEE also outperformed UHGG (67.8% mean mapping rate, Supplementary Figure S1A), demonstrating that local sampling, capturing local diversity, can match or even exceed mapping rates from state-of-the-art global databases built from sample sizes several orders of magnitude larger. Merging of different assemblies led to marginal (i.e., sub-additive) increases in species-level MAGs and negligible improvement in mapping-rates (Supplementary Table S3, Supplementary Figure S1A), implying that most MAGs are redundant (i.e., overlap between multiple datasets).

The large number of species- and subspecies-level MAGs recovered from a moderate sample size suggests that rural Indonesian villages are enriched for microbial diversity and novelty. To quantify this, we compared our newly assembled reference genomes against the largest available reference dataset (UHGG v2.0.1, full dataset: 289,232 MAGs). We discovered that 15% (191) and 49% (1,585) of our species- and subspecies-level MAGs, respectively, did not match the UHGG catalogue (Figure 1B), increasing the number of novel microbial species described from Southeast Asia ∼2.7-fold (Almeida et al., 2021; Gounot et al., 2022). Among novel MAGs, several *Collinsella* species stand out as being both among the most prevalent (up to 43%) and most abundant (0.056%) (Supplementary Table S3). An undescribed *Prevotella* species has the overall highest abundance (0.13%) among novel MAGs and is also moderately prevalent (28%), while a species of the family Anaerovoracaceae is the most prevalent across novel MAGs (45%). While there is no significant difference in abundance overall between novel and known MAGs (Mann-Whitney two-sided test, *p =* 0.15; Figure 1C), novel MAGs are significantly less prevalent (Mann-Whitney two-sided test, *p* < 0.0001; Figure 1D) and are more often found in a single community (Figure 1E, Supplementary Figure S1B) than known MAGs, whilst being distributed widely across the phylogenetic tree (Figure 1F). Further, novel MAGs are found in more similar microbial compositions (i.e., exhibit a lower social niche breadth score (SNB) (von Meijenfeldt et al., 2023)) than known MAGs, as measured by the average pairwise dissimilarity based on the inverse Spearman’s rank correlation (Mann-Whitney two-sided test, *p* < 0.0001; Supplementary Figure S1C-D). The village-specificity of novel MAGs emphasises the importance for local sampling to capture uncharacterised microbial diversity, which here tends to be rare and local. Overall, our novelty results demonstrate unique regional microbiome diversity in Indonesia, which is likely reflective of communities in the broader Southeast Asian region.

### Diverse lifestyle sampling reveals microbiome rural-to-urban transitions

The human microbiome is known to diverge based on geography and lifestyle. In this context, IndoMEE is unique in sampling communities from widely separated tropical islands following diverse subsistence strategies and living conditions, which allows comparison of individuals across a broad socio-cultural and geographic range. We first sought to situate our data through comparison with a nearby, industrial outgroup: Chinese, Malay and Indian groups in Singapore, obtained from the SPMP dataset (Gounot et al., 2022). To better investigate lifestyle variation, we further categorised the analysed communities into remote, rural, and urban groups based on observed parameters, including environment type, food sources, and remoteness (Figure 2A, Supplementary Table S1). Unless otherwise stated, microbiome data was mapped onto the IndoMEE species-level reference database (1,304 MAGs). Controlling for sequencing depth, we find that our Indonesian samples have higher diversity compared to the Singaporean urban outgroup, and identify a gradual and significant decline in diversity from remote to rural to urban communities within Indonesia (Figure 2A; all *p* < 0.05, pairwise Wilcoxon Rank-Sum tests). The decline is consistent across alpha diversity metrics (Shannon diversity index, species richness and Faith’s diversity index), and when samples are mapped under an alternative reference database (merged INDOMEE-UHGG-SPMP database, Supplementary Figure S2A). Within the urban group, Denpasar is significantly more diverse compared to Singapore (*p* = 0.0199, Wilcoxon Rank-Sum test), demonstrating that city-living can be coincident with moderate microbial diversity.

**Figure 2.**
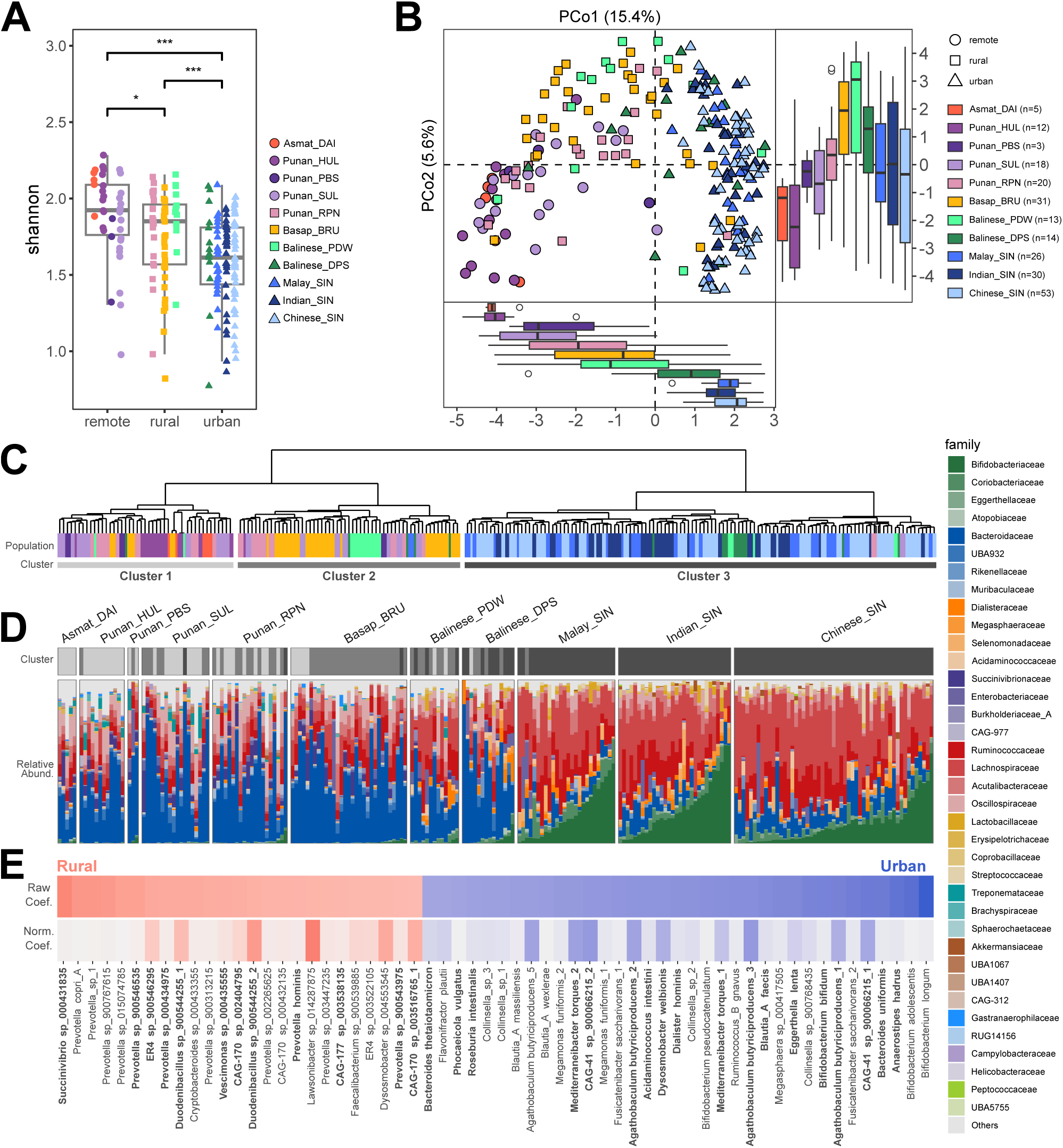
Faecal microbiome diversity across diverse populations spanning a lifestyle cline. **A**) Shannon diversity index for 116 Indonesian and 109 Singaporean individuals across 11 distinct communities. Samples are grouped by lifestyle as remote, rural, and urban. *, ** and *** denote significance thresholds of *p* < 0.05, *p* < 0.01, and *p* < 0.001, respectively. **B)** Principal coordinate analysis of faecal microbiome based on Aitchison distance of 1113 MAGs present in >2 individuals. Box and whiskers plots show the positional distribution of individuals by population along the first and second principal coordinates. Box plot centre lines denote medians, box edges indicate lower and upper quartiles, and whiskers extend to 1.5 times the interquartile range (panels **A** and **B**). **C)** Hierarchical clustering analysis of microbiome beta-diversity distance (Aitchison) groups individuals into three clusters (bottom grayscale bar: remote - light gray; rural - dark gray; urban - black). Node-tips are coloured by population of origin (as in panels **A** and **B**). **D)** Relative abundances of bacterial families across 11 phyla and 11 host communities. Families are ordered by abundance and only the top 4 most abundant families across phyla are shown. The panel above the bar plot shows individuals’ cluster assignment based on panel **C**. **E)** Differential abundance analysis on 561 MAGs presents in >20 individuals in association with the lifestyle cline (ordinal scoring: remote - 0, rural - 0.5, urban - 1), adjusted for age and sex. Eight obese Indonesians were excluded. Only the top 60 associated MAGs by coefficient are shown. The top bar shows the raw coefficients and the bottom bar shows the coefficients normalised by the mean MAG relative abundance. Coefficient heatmaps are coloured by association with the lifestyle cline (rural - red, urban - blue, intensity scaled by the effect size). Species that are also significant in the differential abundance analysis performed on an alternative Indonesia-only dataset are marked in bold.

We further analysed differences in microbiome composition between individuals using principal coordinate analysis (PCoA). Individuals cluster by community, but with a clear trend in variation that reflects the lifestyle cline (from remote to urban) along the first principal axis (15.4% variance explained) (Figure 2B), and divergence of rural villages along the second principal axis (5.6% variance explained). Repeating our PCoA analysis on an Indonesian-only dataset replicated the lifestyle cline (Supplementary Figure S2B), confirming that lifestyle contributes significantly to microbiome composition and alluding to transitional microbiome states as communities become more urban. Hierarchical clustering into three primary sample clusters subsequently assigns samples into groups that are significantly associated with lifestyle categories (Figure 2C; 𝛘^2^ = 225.8; *p* < 0.0001). The influence of lifestyle on microbiome composition is particularly evident in comparisons of geographically-proximate population pairs. Both Bali and Borneo include such pairs, which comprise of communities with divergent lifestyles: remote Punan Tubu Hulu (92% samples in Cluster 1; Figure 2C) shows significantly higher (Shannon) diversity than neighbouring rural Punan Tubu Respen (50% samples in Cluster 2) in Borneo (*p* = 0.0084, one-tailed Wilcoxon Rank-Sum test); and rural Pedawa (62% samples in Cluster 2) shows significantly higher diversity than neighbouring urban Denpasar (64% samples in Cluster 3) despite co-habiting the same relatively small island of Bali (*p* = 0.0434). These results highlight the coherence of a transitional state in the microbiome and the importance of lifestyle over geography in structuring species-level microbiome diversity.

The majority of microbial families (74 of 113, 65%) are significantly different in abundance between sample clusters (Kruskal-Wallis test; *p <* 0.05, Supplementary Table S4). Between urban Cluster 3 and the others (Clusters 1 and 2), the greatest enrichment is observed in Bifidobacteriaceae (44-fold enrichment over Clusters 1 and 2), while the greatest depletions are observed in Treponemataceae (1290-fold), Succinivibrionaceae (64-fold), and Bacteroidales-UBA932 (17-fold) (Figure 2D, Supplementary Table S4), demonstrating broad family-level shifts associated to urbanisation. To explore differential abundance at the species-level, we used MaAsLin2 (Mallick *et al.,* 2021), focussing on 561 MAGs with prevalence > 20 individuals. We find 364 (67%) MAGs significantly associated with the rural-to-urban lifestyle cline, emphasising the breadth and scale of microbial response (*p* < 0.05, adjusted for age and sex, Supplementary Table S4). Many associations (151/364, 41%) are replicated in an Indonesia-only analysis, despite lower sample size and power, with coefficients consistently in the same direction (Supplementary Table S4). Notably, four *Bifidobacterium spp.* (*B. longum. B. adolecentis, B. bifidum,* and *B. pseudocatenulatum*) are among the most associated species with urbanisation in the Indonesia-Singapore dataset (Figure 2E). We find that *Bifidobacterium spp.* enrichment is negatively correlated with *Treponema succinifaciens* (Pairwise Kendall 𝛕 = −0.20 to −0.30, all *p* < 0.01), in accordance with previous work on rural and hunter-gatherer microbiomes (Angelakis et al., 2019; Rosas-Plaza et al., 2022; Schnorr et al., 2014; Stražar et al., 2021). Notably, among *Bifidobacterium* species, only *B. bifidum* is significantly associated with urbanisation in the Indonesia-only dataset (Figure 2E, Supplementary Table S4). *B. bifidum* occurs only in Denpasar, Basap, and Singapore, while other *Bifidobacterium* species are present in the vast majority of samples at 27‒43 fold lower abundance in Indonesia than Singapore. We also observe generally reduced abundance of *Prevotella (P. copri, P. hominis,* and other *Prevotella* species) and increased abundance of *Bacteriodes/Phocaeicola* (e.g., *B. uniformis*, *B. thetaiotaomicron*, and *P. vulgatus*) with increasing urbanisation (Supplementary Table S4). Other notable enrichments in the urban populations include Bacillota A members: *Anaerostipes hadrus, Agathobaculum butyriciproducens, Blautia_A faecis, Blautia_A wexlerae*, and *Ruminococcus_B gnavus*; along with Actinomycetota members: *Eggerthella lenta* and *Collinsella sp.*; while Pseudomonadota members: *Succinivibrio sp.* and *Duodenibacillus sp.* are depleted. Indeed, clines in relative abundance over the lifestyle spectrum are observed for a wide range of genera and species (Supplementary Figure S2C).

### Community, more than individual, lifestyle predicts microbiome differentiation in Indonesia

To further investigate the drivers of this lifestyle signal, we analysed individual-level diet and daily activity data available for 122 Indonesian samples from 5 of the sampled communities (Supplementary Figure S3A-B). This comprised consumption frequency across 7 core food categories (rice, sago, pork, egg, milk, fish and chicken) and 2 daily activities (hunting and watching television) (see Methods and Supplementary Table S2). Principal component analysis (PCA) of this expanded lifestyle data reveals substantial variation, highly structured by community, with PC1 (26% variance explained) driven by hunting as well as consumption of sago (palm) and pork (wild in remote/rural communities) in Asmat and Punan Tubu Hulu (Figure 3A, Supplementary Figure S3C). These individuals contrast those from Pedawa and Denpasar, who had a higher frequency of watching television and chicken consumption. On the other hand, PC2 (15.7% variance explained) is driven by higher consumption frequencies of fish, egg and milk (Figure 3A, Supplementary Figure S3C).

**Figure 3.**
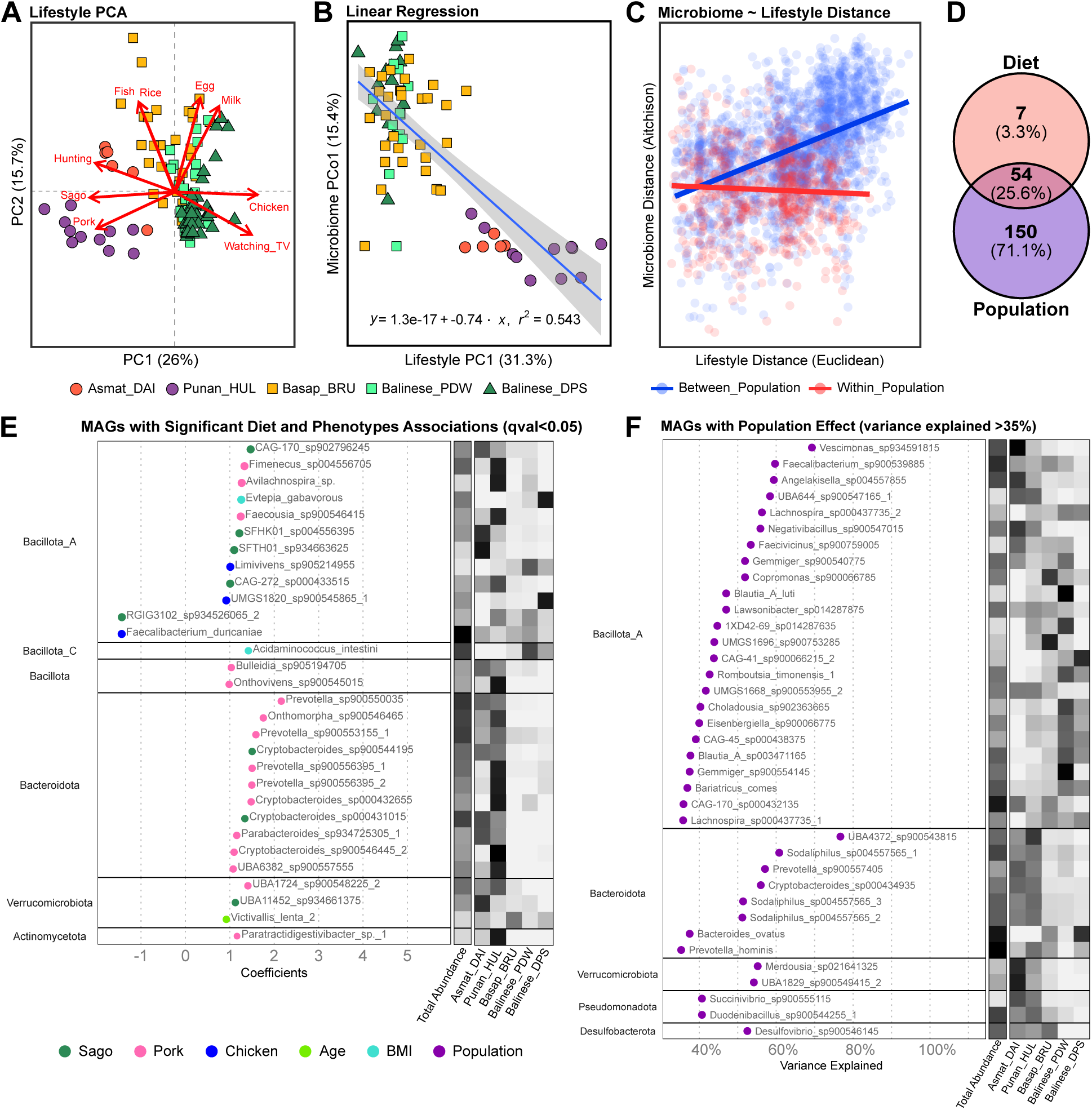
Microbiome composition is associated with lifestyle factors in Indonesia. **A)** Principal component analysis (PCA) of lifestyle variables from 122 individuals across five Indonesian populations (communities). Samples are coloured by population (see legend below panel). **B)** Linear regression between the first principal component (PC1) of the lifestyle PCA and PCo1 of the microbiome PCoA, based on 73 individuals with complete metagenomic and lifestyle data. Samples are coloured by population (see legend below panel). **C)** Pairwise relationships between microbiome distances (Aitchison) and lifestyle distances (Euclidean), stratified by between-population (1,941 pairs, blue; Mantel test ρ = 0.42, *p* = 0.0001) and within-population (687 pairs, red; Mantel test ρ = -0.04, *p* = 0.6528) comparisons. **D)** Venn diagram comparing the number of MAGs significantly associated (*p* < 0.05) with diet and population, as identified by MaAsLin2. **E)** MAGs significantly associated with diet or phenotype (age, sex, and BMI) under a full mixed effects model with population as a random effect (*p* < 0.05). MAGs are coloured by their association with sago, pork, chicken, age, or BMI (see legend below panel). The heatmap panel shows total and normalised mean abundance across the five populations. **F)** MAGs that are strongly associated with population (>35% of variance explained by population), under the same mixed effects model of panel E.

We observe a significant correlation between the first principal axes of the microbiome PCoA and lifestyle PCA (R^2^ = 0.543, *p <* 0.0001; Figure 3B), with notable divergence of remote groups from others. Under a mixed model with population as a random effect, however, this association is not significant (*p* = 0.1026; 57.8% variance explained by population), suggesting that lifestyle differences between populations may underlie this relationship. Indeed, we do not observe significant correlations between the first principal axes of lifestyle and microbiome ordinations within individual groups (Supplementary Figure S3D). Moreover, correlation of microbiome and lifestyle distance between sample pairs (Figure 3C) reveals a significant correlation between populations (Mantel test, ρ = 0.42, *p* < 0.0001) but not within population (Mantel test, ρ = −0.04, *p* = 0.6528). Across population pairs, we find a significant correlation between microbiome divergence even when controlling for geography (partial Mantel test, ρ = 0.57, *p* = 0.0417; Supplementary Figure S3E), indicating that between-community divergence is not purely a spatial effect. Indeed, no such significant correlation is found when correlating microbiome divergence with geographic distance controlling for lifestyle difference (partial Mantel test, ρ = 0.18, *p* = 0.3250). Together, our results offer compelling evidence that microbiome divergence is driven by lifestyle differences structured at the scale of communities.

We then explored if MAG relative abundance could be predicted by specific diet components. Variable selection based on univariate associations with seven core diet variables (rice, sago, pork, egg, milk, fish, and chicken) identified three primary predictors: sago, pork and chicken. To compare the contribution of diet and population to MAG abundance, we analysed two models in MaAsLin2: a multivariate diet model (MAG_abundance_ ∼ sago + pork + chicken + age + sex + BMI) and a population model (MAG_abundance_ ∼ population + age + sex + BMI); with age, sex and BMI included to control for potential confounding effects. Across 667 MAGs tested, 61 (9.14%) show significant dietary associations, while 204 (30.58%) are significantly associated with population; notably, the vast majority (54/61, 88.52%) of dietary associations are also detected as population-associated (Figure 3D, Supplementary Figure S3F). To better account for variable interaction and identify true diet associations, we then used a mixed effects model incorporating the three dietary components as fixed effects and population as a random effect (MAG_abundance_ ∼ sago + pork + chicken + age + sex + BMI + 1|population). This yielded 27 (4.05%) diet-associated MAGs (Figure 3E), most of which are associated with pork consumption. Population also accounted for a considerable proportion of variance explained (>35%) in 37 MAGs (Figure 3F). MAGs most influenced by population include Prevotellaceae bacteria *UBA4372 sp900543815* (76.7% variance explained), *Vescimonas sp.* (69.3%), *Sodaliphilus sp.* (60.8%), *Faecalibacterium sp004557565* (59.7%), and *Angelakisella sp004557855* (59.4%). In contrast, the variance explained by population in diet-associated MAGs ranged between 0‒18.7%. Taken together, our analyses incorporating lifestyle data emphasise that host population is a stronger predictor of microbiome composition than individual diet, likely reflecting population-scale properties such as shared community lifestyle (including diet, see Figure 3C), environment, and microbial transmission.

### Sporulation as a key bacterial trait that cascades up to the structure and novelty of the social microbiome

The composition of the gut microbiome in Indonesian populations is predicted in part by host lifestyle and community structure, but may also be driven by microbial traits that influence transmission and survival in non-host environments. We functionally annotated MAGs across a broad suite of functional classes including carbohydrate-active enzymes (CAZy), KEGG functional orthologs, anti-microbial resistance (AMR) genes and others (see Methods). We identify functions across the spectrum of the aforementioned classes, and a large number of uncharacterised (427,699) and novel (116,953) proteins, in our dataset (2,129,118 proteins total). Novel MAGs are enriched for both uncharacterised and novel proteins relative to known MAGs (𝛘^2^ tests, all *p* < 0.0001). Focusing on KEGG functional orthologs (n = 6,450), carbohydrate-active enzymes (n = 325), AMR genes (n = 115) and others (Supplementary Figure S4), we performed PCA and find that clines in species composition (Figure 2B) are mirrored by functional structure, with ca. 20‒27% variance explained by PC1, and with consequent strong association with the lifestyle cline (Supplementary Figure S4).

We then assessed which (KEGG) functions or genes are enriched or depleted in the novel MAGs relative to the known MAGs. Of 157 genes identified as significantly enriched or depleted, a significant proportion (17%; 27 genes) are identified as associated with sporulation (Figure 4A). Notably, all of these are inferred as significantly depleted in the novel MAGs. The transmission of microbes depends in part on their persistence in non-host environments, which can be facilitated by the development of a spore that exhibits minimal metabolic activity and high resistance to environmental stressors (Swick et al., 2016). However, a link to novelty has been less clear, though it may be hypothesised that this association (i.e., between novelty and sporulation) could be driven by lower prevalence (Browne et al., 2021). To test if depletion in sporulation genes is associated with lower prevalence (and consequently MAG discovery), we regressed MAG sporulation gene counts *n*_s_ (i.e., the number of distinct sporulation-associated genes present in a MAG; Supplementary Figure S5A) against MAG prevalence and the number of villages the MAG is present in. As bacterial spore-formation is a characteristic trait of Bacillota (previously Firmicutes) (Galperin, 2013), we assessed this both within and across Bacillota phyla, in addition to across the whole dataset (i.e., all MAGs across all phyla). Higher *n*_s_ is significantly associated with higher prevalence in the whole, pan-Bacillota and Bacillota A datasets, while results are insignificant (or significant in only one of the prevalence metrics) for Bacillota B, Bacillota C and Bacillota ss (Figure 4B, Supplementary Figure S5B-E). This may be partly attributed to the low sample size of these latter phyla compared to Bacillota A in our dataset (Figure 1F). Results across various annotations (e.g., KEGG and Pfam) are consistent, suggesting results are robust to annotation database (Supplementary Figure S5B-E). We observe that prevalence cascades to the village-level, with non-spore-forming bacteria more often unique to a single village than spore-forming bacteria, which are more ubiquitously shared across villages (two-sample Kolmogorov-Smirnov test, D = 0.17, *p* < 0.0001; Figure 4C). Additionally, we note that the master regulator gene essential for sporulation, Spo0A, is a strong predictor of *n*_s_ (*p* < 0.0001; Supplementary Figure S5F), further supporting the link between sporulation and prevalence.

**Figure 4.**
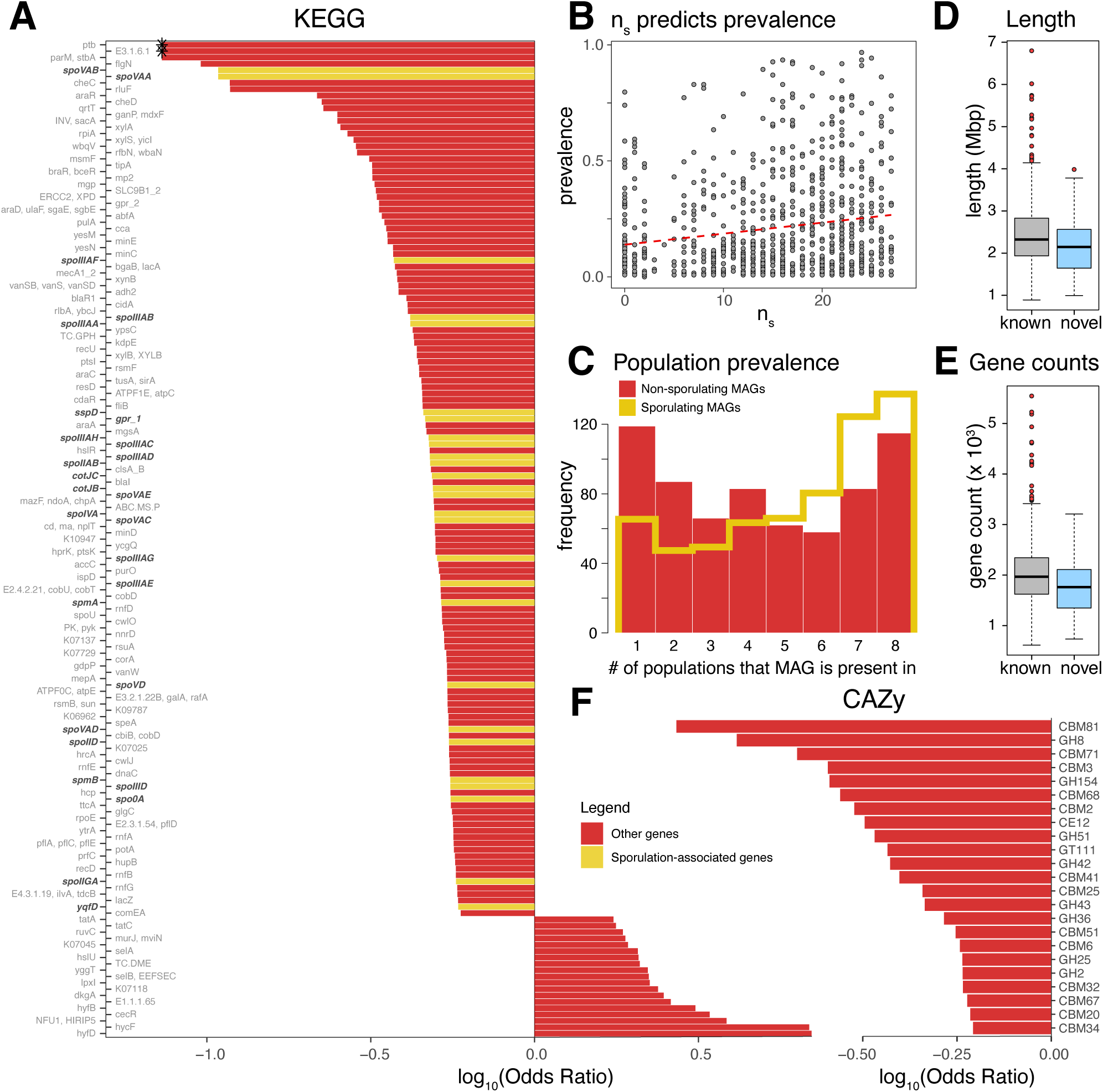
Sporulation gene counts, which are depleted in novel MAGs, predict microbe prevalence. **A)** Functional enrichment analysis (KEGG functional orthologs) of novel versus known MAGs, showing all significant enriched and depleted features. Sporulation-associated genes are coloured in yellow and annotated in bold. Significance and odds ratios were calculated via Fisher’s exact test, with negative odd ratios indicating depletion in novel MAGs. Asterisks denote infinite odds ratios (i.e., zero denominator). **B)** MAG sporulation gene count (*n*s) is associated with higher prevalence across Bacillota phyla (linear regression; *p* < 0.0001). **C)** Non-sporulating MAGs (red histogram) are more often unique to a single village than sporulating MAGs (yellow outlined histogram), which are more ubiquitously shared across villages (two-sample Kolmogorov-Smirnov test, D = 0.17, *p* < 0.0001). Here, sporulating or non-sporulating is discriminated by the presence or absence of the master regulator gene essential for sporulation, Spo0A. **D)** Length of novel MAGs compared to known MAGs. Novel MAGs are significantly smaller than known MAGs (Mann-Whitney one-sided test, *p =* 0.0014). **E)** Gene count of novel MAGs compared to known MAGs. Novel MAGs possess significantly fewer genes than known MAGs (Mann-Whitney one-sided test, *p =* 0.0032). In **D**) and **E**), centre lines denote medians, box edges indicate lower and upper quartiles, and whiskers extend to 1.5 times the interquartile range. **F)** Functional enrichment analysis of novel versus known MAGs for carbohydrate-active enzymes (CAZy), showing all significant features. Significance and odds ratios were calculated via Fisher’s exact test, with negative odd ratios indicating depletion in novel MAGs.

Loss of sporulation capacity in Bacillota has been associated with greater host adaptation, and has been found to coincide with a suite of contemporaneous adaptations, including reduced genome size and narrowed metabolic capabilities (Browne et al., 2021; Debray et al., 2024). Consistent with these expectations, we find that novel MAGs in IndoMEE are smaller (Figure 4D; Mann-Whitney one-sided test, *p =* 0.0014), possess fewer genes (Figure 4E; Mann-Whitney one-sided test, *p* < 0.0001) and exhibit reduced metabolic carbohydrate metabolism (Figure 4F) compared to known MAGs, implying that novel MAGs may be particularly enriched for host-adapted species.

Collectively, our results elucidate links between bacterial traits (i.e., sporulation capacity), degree of host adaptation, transmission and, ultimately, prevalence; with reduced prevalence consequently impacting the probability of previous sampling in existing databases and hence novelty in our analysis. Sporulation hence appears to be a key bacterial trait that cascades up to the structure and novelty of the social microbiome, and explains in part the village-level differentiation patterns identified across rural Indonesia.

## Discussion

Our work presents a new Indonesian gut microbiome dataset, IndoMEE, that incorporates diverse lifestyle sampling and substantially extends known MAG diversity of the Southeast Asian region. We find 15% of species-level MAGs and almost 50% of subspecies-level MAGs are novel, emphasising the limited representation of both non-industrial populations generally and Southeast Asia in existing databases. Intriguingly, many novel species have low-prevalence, often found in only a single village. These novel species show depletion of sporulation genes, and indeed we find a clear overall association between sporulation gene count and prevalence – demonstrating the impact of microbial traits cascading up, likely via transmission efficiency, to influence the distribution of taxa, and ultimately species discovery. The varied lifestyles represented by our sampling allowed us to study lifestyle-microbiome associations in detail. We identify key differences between Indonesian and highly urbanised Singaporean samples, and observe clines in abundance among a wide range of taxa. Within Indonesia, we find that an individual’s community-membership is a key predictor of microbiome composition. While diet has a pronounced impact on microbiome composition, our results suggest that this is mediated by and structured at the scale of communities; an important observation for understanding human gut microbiome variation.

Lifestyle transitions coincide with significant changes in diet (Dounias et al., 2007; Popkin, 2004, 2011), hygiene practices (Dounias & Froment, 2011), healthcare (including antibiotic use), resource acquisition strategies (Reyes-García et al., 2019), and human-environment interactions, and have been associated with significant changes in the gut microbiome. For example, urban populations show a characteristic reduction in gut microbiome diversity compared to various agricultural and hunter-gatherer communities, in addition to compositional differences such as decreased *Prevotella* abundance (Mancabelli et al., 2017; Rosas-Plaza et al., 2022; Sonnenburg & Sonnenburg, 2019; Vangay et al., 2018). How general or varied compositional trends are across different populations and lifestyles, however, remains unclear, given the continued limited representation of non-Western, non-industrialised microbiome samples. Our work reveals intermediate (transitional) states in microbiome composition along our sampled lifestyle cline that spans hunter-gatherer, agricultural and urban contexts in Indonesia. Specific community comparisons are especially informative when viewed in historical context, given that three Bornean communities (Punan Tubu Respen, Punan Aput and Basap) transitioned from traditional hunter-gatherer to settled agricultural lifestyles in the 1970s/1980s (Zahorka, 2017), and given rapid and recent economic development in Bali. For example, while urban Bali Denpasar clusters with Singapore, their microbiomes are also compositionally similar to more rural Indonesian groups; microbiome samples from the long-term agricultural Bali Pedawa and recently transitioned Punan Tubu Respen communities have similar composition despite different subsistence histories and disparate geographies (i.e., Bali and Borneo); and Punan Tubu Respen and Punan Tubu Hulu, who are from the same ethnic community but integrated into the market-economy to different extents over recent generations, have distinct microbiome compositions. Overall, our work complements studies of inter-generational shifts among migrants (Vangay et al., 2018) and supports relatively rapid changes in community microbiomes ‒ over the timescale of a few generations ‒ in ways that are strongly predicted by lifestyle.

Our described lifestyle cline is further evident at the level of microbial taxa. For example, we find enrichment of *Bifidobacterium* species and depletion in *Treponema* species in urban groups, as has been previously reported (Angelakis et al., 2019; Gounot et al., 2022; Jha et al., 2018; Nakayama et al., 2015; Stražar et al., 2021). *Bifidobacterium* are prevalent in the human gut, particularly during infancy (Derrien et al., 2022), and the association with urbanisation may reflect increased accessibility to wheat, dairy, and probiotic products (Chonnacháin et al., 2024; Lu et al., 2023), with potential immune response implications (Stražar et al., 2021). We also identify a *Prevotella* and *Bacteroides* association with lifestyle, as has been previously described (Gorvitovskaia et al., 2016). While *Bacteroides* and *Prevotella* both metabolise polysaccharides, they differ in bile tolerance, and so increased intake of animal fat and meat as associated with rural-to-urban dietary shifts is thought to favour *Bacteroides*. While hunter-gatherer and rural diets vary, enrichment of *Prevotella* has been observed in several hunter-gatherer communities (Rosas-Plaza et al., 2022). We also identify a range of new lifestyle-associated taxa ‒ indeed, more than half (67%) of MAGs tested are associated with the remote-rural-urban axis, with pronounced signals across a wide range of genera. Our study extends previous work on microbiome-lifestyle associations in Indonesia based on 16S data (Febinia et al., 2022; Nakayama et al., 2015), and highlights transitional states of the microbiome in a rural-to-urban lifestyle cline.

Major shifts in the microbiome can occur rapidly at the individual scale (David et al., 2014), however, the nature of lifestyle-associated microbiome transitions in human communities is less clear. Our results demonstrate that lifestyle associations are driven more by between-community than within-community differences. Diet remains a fundamental driver of microbiome composition, but may explain a minor part of compositional variance as noted across a range of study designs (Johnson et al., 2020). Our results thus emphasise the importance of population-level processes – e.g., shared lifestyle (including diet), environments and more active processes like food sharing and transmission (Debray et al., 2024; Valles-Colomer et al., 2023) – in shaping the social microbiome of human communities.

One limitation of this study is that the lifestyle differences we analyse may be confounded by demographic and methodological confounders. This concern led us to replicate key results using relevant subsets of the overall dataset, namely an Indonesia-only analysis of the lifestyle cline (Figure 2E, Supplementary Table S4), and an Indonesia (excluding Balinese) analysis of diet and population MAG associations (Supplementary Figure S3F). Assessment of the effect of confounders on microbiome composition through PERMANOVA model selection (see Methods) identified study protocol as the only relevant confounder, with lifestyle nevertheless retained as the most predictive variable (Supplementary Table S4). Replication and statistical results, along with alignment of findings with key prior expectations (such as reduced diversity in urban contexts, or increased representation of *Bifidobacterium* in cities), collectively support a robust association of lifestyle with microbiome composition. Our results, particularly related to diet associations, are also contingent on the representativeness of our diet data that was collected via a food frequency questionnaire (see Methods). While we strived to incorporate all staple and frequently consumed foods to capture key dietary variation within and among sampled communities, this data does not reflect, nor does it intend to evaluate, the entirety of an individual’s nutritional profile, which would necessitate other forms of assessment, such as meal frequency, caloric intake, and nutritional density.

Beyond lifestyle, our work points towards the role of microbial traits, specifically sporulation, in revealing host-microbe dependence and in shaping microbiome structure. In our study, novel MAGs are significantly depleted for sporulation genes. Sporulation in bacteria is exclusive to Bacillota (previously Firmicutes) (Galperin, 2013), where sporulation is considered an ancestral trait. A loss of sporulation capacity has been linked to greater host adaptation, as it mandates more direct modes of between-host transmission to avoid non-host environmental exposure (Browne et al., 2021; Debray et al., 2024). The resultant, altered transmission cycle and greater reliance on host are hence expected to drive a suite of contemporaneous adaptations, including reduced genome size and narrowed metabolic capabilities (Browne et al., 2021). Our novel MAGs exhibit these tell-tale signatures of host dependence, suggesting that they may be particularly enriched for host-adapted species, and suggesting a mode of transmission via interpersonal contact more so than environmental dispersal. Indeed, we reveal that microbe sporulation capacity correlates significantly with prevalence (Figure 4, Supplementary Figure S4G-H, S5). While a link between sporulation, transmission and prevalence has been proposed (Browne et al., 2021), the broader implications of this for characterising microbiome diversity and particularly MAG discovery has remained unclear. Our results draw a direct link between microbial physiology, prevalence, the social microbiome, and MAG discovery.

Our diverse gut microbiome sampling from Indonesia substantially extends known microbial genome diversity in Southeast Asia, and identifies clinal patterns of microbiome variation that mirror community-scale lifestyle trends. By combining analyses on novelty and function with prevalence and population structure, our work further highlights a conceptual bridge between bacterial physiology, prevalence, community divergence and species discovery. Further sampling efforts in under-represented regions like Indonesia, particularly at the village scale and spanning diverse lifestyles, will likely discover many more low-prevalence, often non-sporulating and host-adapted species, and offer further insights into the processes structuring microbiome composition at local and global scales.

## List of supplemental information

Supplementary Information document. Figures S1–S5 and Table S1.

Table S2. Excel file containing additional data too large to fit in a PDF, related to Figures 1-3.

Table S3. Excel file containing additional data too large to fit in a PDF, related to Figure 1.

Table S4. Excel file containing additional data too large to fit in a PDF, related to Figures 2-3.

## Data and code availability

Raw metagenomic sequencing reads for the 116 faecal samples are deposited at the European Genome-phenome Archive (EGA), under accession number EGAS50000000961. IndoMEE MAGs are available on Figshare at https://doi.org/10.6084/m9.figshare.28660445. Code used in this study is available at the GitHub repository: https://github.com/hirzi/IndoMEE.

## Acknowledgements

The authors thank communities and volunteers involved in this research, as well as Prof. J Stephen Lansing, A/Prof. Andrew J. Holmes, Prof. I Wayan Weta, Datuk Karim, Ahmad Arif, members of the Genome Diversity and Diseases Laboratory field team (Prisca C. Limardi, Isabella Apriyana, Gludhug A. Purnomo, Bertha L. Utami, and Leonard Taufik), and university students at the Faculty of Medicine of Udayana University for supporting and assisting the community engagement and sample collection of this study between 2015 and 2021. We are grateful to local officials in Tanjung Selor and Malinau, and the directors of the Eijkman Institute for Molecular Biology (Prof. Sangkot Marzuki and Prof. Amin Soebandrio) for facilitating this study. This research was funded by the European Union’s Horizon 2020 research and innovation programme under grant agreement no. 950610 and an NTU Presidential Postdoctoral Fellowship to G.S.J. (2018-2020), a Wellcome Trust International Training Fellowship (no. 222992/Z/21/Z) to P.K., a block grant from the government of the Republic of Indonesia to the Eijkman Institute for Molecular Biology through the Ministry of Research and Technology/National Agency for Research and Innovation (IMELDA Project, 2014‒ 2021), the International Program Development Fund to A/Prof. Andrew J. Holmes, H.S., and S.G.M. (2013), Australia’s Department of Foreign Affairs and Trade through the Australia Awards Scholarship Fieldwork Entitlement to C.A.F. (2014‒2016), and the Cambridge Trust Scholarship to C.A.F. (2022‒2024).

## Author’s contributions

C.A.F., H.L., S.G.M, and G.S.J. conceptualised the study. C.A.F., P.K., L.P., D.W., G.I., H.S. and S.G.M. facilitated and performed sample and data collection. C.A.F., H.L., P.K. and L.P. performed data curation. C.A.F., P.K., L.P., S.G.M., and G.S.J. developed field and laboratory protocol. C.A.F., H.L., A.A., and G.S.J. developed bioinformatic and statistical analysis pipelines. H.L. performed metagenomic assembly and annotation; and C.A.F., H.L., and J.L. performed data analyses. C.A.F., H.L., and G.S.J. wrote the manuscript, which all authors revised. G.S.J. and S.G.M. administered and supervised the study.

## Methods

### Study enrolment

Community engagement and sample collection started in 2015 and was completed in 2021. Prior to collecting the samples, we liaised with local officials and leaders (e.g., village elders) to explain and seek permission to perform the study. During enrolment, we collected written informed consent from the volunteers, following community meetings that introduced the study. Data and sample collection were done in one instance (cross-section) in each community on separate occasions: Balinese Denpasar (Udayana University) in 2015, Punan Tubu Respen in 2018, Punan Aput in 2018, Basap in 2019, Punan Batu in 2019, Asmat Daikot in 2019, Punan Tubu Hulu in 2020, and Balinese Pedawa in 2021 (Supplementary Table S1). We included only adults in this study (≥18 years old), and additionally excluded i) individuals with recent gastrointestinal symptoms such as diarrhoea (<3 days), ii) individuals with a close kin relationship (siblings and parents), iii) pregnant women, and iv) individuals with chronic liver and kidney diseases. We were unable to firmly exclude all individuals with recent antibiotic use given the limited availability of medical records, and challenges in confidently explaining the difference between antibiotics and other medicines to community members (participants).

The study was approved by the Eijkman Institute for Molecular Biology Research Ethics Commission (project No. 80/2014, No. 119/2019, and No. 126/2019), the Udayana University Faculty of Medicine and Sanglah Hospital Ethics Commission (No. 1286/UN.14.2/Litbang/2014 and No. 1742/UN14.2.2.VII.14/LT/2021), the Mochtar Riady Institute for Nanotechnology Ethics Committee (006/MRIN-EC/ECL/III/2022), and the University of Cambridge Human Biology Ethics Committee (HBREC.2021.01).

### Data and sample collection

We collected participant metadata including age and family tree. We assessed dietary intake and daily activities (i.e., hunting, farming, walking) through interviews. In total, we interviewed 172 individuals for lifestyle data and obtained complete dietary data from 122 individuals, of which 12 were identified as obese (body mass index (BMI) > 30).

The diet interview was conducted using a semi-quantitative food questionnaire (FFQ), in which participants reported the number of servings of specific food items (e.g., white rice) consumed and the frequency (daily, weekly, monthly, or yearly). The questionnaire included a list of commonly consumed foods, mainly staple carbohydrates (e.g., rice, sago, tubers, potatoes, and noodles), meat products (e.g., chicken, pork, fish, and beef), legumes, leafy vegetables, eggs, dairy products, and other market products (e.g., snacks and confectioneries). In subsequent analyses, the diet data was reduced to a common core set of seven food categories (rice, sago, pork, egg, milk, fish, and chicken), and converted to yearly frequency. This FFQ was developed in collaboration with a local clinical nutritionist and a public health expert (see Acknowledgements, Febinia et al., 2022). We note that our FFQ was specifically designed to measure commonly consumed food items. It does not measure other dietary aspects (e.g. nutrition density, caloric intake) and therefore it is not intended as a general nutritional assessment. We also note that as a self-reported instrument, it remains subject to known limitations, such as recall bias and self-reporting errors (Cade et al., 2004).

The physical activity data captured the intensity of routine activities by measuring their frequency (times per day, week or month) and duration (in hours or distance). Examples included farming, hunting, foraging, fishing, and commuting (whether on foot or by other means). Questions on sedentary behaviours such as sleeping, watching television, and casual socialising were also included. Use of vehicles was recorded if relevant.

Faecal samples were obtained from 116 individuals across the eight communities, of which 73 had fully completed the lifestyle questionnaire. For Bali (Pedawa and Denpasar), faecal samples were stored in −20 °C within 4 hours of defaecation and subsequently transferred to −80 °C upon arrival at the lab. For other communities, due to limited access to cold storage, samples were preserved at room temperature using OmnigeneGUT (DNA Genotek, Cat# OMR-200) for a maximum of 60 days, adhering to the manufacturer’s guidelines. Prior work has found limited differences between these storage methods (Guan et al., 2021; Ilett et al., 2019; Song et al., 2016); nevertheless, we consider this batch effect in our analyses. Singaporean metagenomic shotgun sequence data (derived from faecal samples) were obtained from the public repository of the Singapore Platinum Metagenomes Project (NCBI project accession number PRJEB49168) (Gounot et al., 2022).

### Lifestyle categories

In this study, we categorised the sampled communities into remote, rural, and urban groups to facilitate comparisons among populations with shared traits. This categorisation is based on observed parameters, including environment type, food sources, and remoteness (detailed in Supplementary Table S1). Specifically, Asmat Daikot, Punan Batu, Punan Tubu Hulu, and Punan Aput are classified as remote; Punan Tubu Respen, Basap, and Balinese Pedawa as rural; and Balinese Denpasar and the Singaporean samples as urban. These classifications are not intended as official designations; acknowledging the variability in definitions across regions and countries, and the interplay of factors such as population density, proximity to economic centres, infrastructure, economic diversity, and development across sectors.

### DNA extraction and sequencing

DNA extraction was performed within three weeks of sample collection to minimise storage time. Extraction was conducted in a Level-2 Biosafety Cabinet using the PowerFecal DNA Isolation Kit (MO BIO, Cat# 12830-50) for the Denpasar samples. For the remaining samples, the QIAamp PowerFecal DNA Kit (QIAGEN, Cat# 2830-50), the rebranded version of the original kit, was used. Extractions used 0.25 g of stool for each sample. We included extraction blanks (250 μL elution buffer 10 mM Tris, Qiagen Cat# 19086) as negative controls. For the Denpasar samples, homogenisation was performed using a Vortex Genie II machine (Sigma-Aldrich, Cat# Z258423) at maximum speed for 15 minutes (Febinia et al., 2022). The remaining samples were homogenised using the FastPrep-24 system (MP Biomedicals, Cat#116004500) at 5.5 m/s for two cycles of 30 seconds each, with a 5-minute interval between cycles. The quantity of extracted DNA was measured using the Qubit dsDNA Broad Range Kit (ThermoFisher Scientific, Cat# Q32853). DNA purity was assessed with a Nanodrop Spectrophotometer ND-1000 (ThermoScientific), and genome integrity was evaluated via gel electrophoresis in 1% agarose using 1× TBE buffer. Samples with contamination or low DNA quantity underwent a second (repeat) extraction. If the quality improved, the second extracts were used for further analysis.

We sequenced a total of 116 samples in total, in addition to 8 technical replicates, 2 positive controls (ZymoBIOMICS Microbial Community DNA Standard by Zymo Research Cat# D6305), and 4 negative controls (for sampling: blank, for extraction: Buffer EB by Qiagen Cat# 19086, for library prep: Nuclease Free Water by Ambion-Invitrogen Cat# AM9937). DNA libraries were prepared using Illumina’s TruSeq Nano DNA Kit (Illumina, Cat# 20015965. Sequencing was performed on the Illumina NovaSeq 6000 platform to generate 250 bp paired-end reads, with an average yield of 10.63 Gb / sample. Following quality control, we acquired high-quality faecal metagenomic sequences from 116 samples for subsequent analyses (Supplementary Table S2).

### Metagenomic assembly

We assembled metagenome-assembled genomes (MAGs) using a modified version of the metaWRAP pipeline (version 1.3.2) (Uritskiy et al., 2018), from the metagenomic sequence data of 116 samples. Reads were first trimmed for adapters and by quality using Trim Galore (version 0.6.10) (Krueger, 2016), under default settings (-q 20 --length 20 -e 0.1). Trimmed reads were then aligned to the complete human (host) reference genome (CHM13v2.0) via bmtagger (version 1.1.0), to remove human reads from the metagenomic sequences.

Host-cleaned metagenomic reads were then assembled via a mixed approach that combined individual-and co-assembly methods. Combining reads via co-assembly increases read depth which facilitates the recovery of species that may be too rare to detect in individual samples, however, mixing data from numerous strains can adversely affect the assembly process, as intraspecific genomic variation can cause de Bruijn graphs to incorporate numerous alternative paths (Delgado & Andersson, 2022). Here, co-assembly was performed per population, i.e., metagenomic reads of individuals were pooled within population (not across populations), resulting in 7 population FASTQ files. Reads were assembled using MEGAHIT (version 1.2.9) for each individual (individual assembly) and for each population (co-assembly). Assembled contigs were then binned via MaxBin2 (version 2.2.4), metaBAT2 (version 2.12.1) and CONCOCT (version 1.0.0) (individual assembly) or MaxBin2 exclusively (co-assembly). When more than one binning software was used (individual assembly), we used metaWRAP’s bin-refinement module to produce a consolidated, improved MAG set, based on MAG completeness and contamination rates as estimated from CheckM (version 1.0.18) (Parks et al., 2015). Specifically, MAGs were scored based on the scoring function *S*_1_ *= Completeness – 5 * Contamination*. We additionally applied a hard filter, requiring a minimum completeness of 50% and allowing a maximum contamination of 5% for MAGs. This yielded 8,633 and 2,437 MAGs for the individual assembly and co-assembly respectively. The resultant set of MAGs are non-redundant and comprise many replicates across individuals and across populations. To get representatives at species and subspecies levels, MAGs were dereplicated using dRep (version 3.4.5) (Olm et al., 2017) at 95% and 98% average nucleotide identity (ANI) respectively, using the fastani algorithm (Jain et al., 2018), CheckM2 version 1.0.2 (Chklovski et al., 2023) estimates of completeness and contamination, and the following dRep parameters: -pa 0.9 -sa 0.95 -nc 0.3 -l 50000 -comp 50 -con 5. The genome with the highest *S*_2_ score was chosen as the cluster representative, where *S*_2_ *= A * Completeness - B * Contamination + C * (Contamination * (strainHeterogeneity / 100)) + D * log(N50) + E * log(size) + F * (centrality - Sani)*, and A-F are coefficients with values 1, 5, 1, 0.5, 0 and 1, respectively (Olm et al., 2017). Dereplicated MAGs were subsequently filtered for *S*_1_ ≥ 50% and for chimerism and contamination content via GUNC (version 1.0.6) (Orakov et al., 2021). For the latter, we filtered out genomes that matched all the following criteria: i) flagged by GUNC (i.e., clade_separation_score > 0.45; contamination_portion > 0.05; reference_representation_score > 0.5), ii) are singletons (dRep clusters with only one member) and iii) < 90% completeness based on CheckM2. Finally, we combined and dereplicated the resultant individual assembly and co-assembly MAGs, using the same dRep parameters as above, to generate the final assembly.

### Taxonomic assignment and phylogenetic tree construction

To taxonomically classify MAGs, we used GTDB-tk (version v2.3.2) (classify_wf) to assign taxonomies to MAGs based on the Genome Taxonomy Database (GTDB release 214), and to build multiple sequence alignments for constructing phylogenetic trees. We generated phylogenetic trees via IQ-TREE (version 2.3.3) (Nguyen et al., 2015) with the ModelFinder Plus (-m MFP) argument, which automatically performs model selection and subsequently constructs trees using the inferred best-fit model. Best-fit models according to the Bayesian information criterion (BIC) for bacteria and archaea were LG+R10 and LG+F+G4, respectively. We then used the Interactive Tree Of Life (iTOL) (version 6) online plotting tool to annotate and plot phylogenetic trees.

### Quantifying abundance, prevalence and mapping rates

To evaluate mapping rates across different assemblies, and calculate MAG relative abundances, we used metamap (Almeida, 2021c). metamap maps paired-end metagenomic reads to a reference genome using bwa-mem (version 0.7.17-r1188) (Jung & Han, 2022; Li, 2013), and filters mapped reads based on nucleotide identity, mapping score and alignment fraction using SAMtools (version 1.9) (Danecek et al., 2021). For each MAG in the reference, read counts, depth and (observed and expected) breadth of coverage are then calculated; with expected breadth given as *breadth_exp_ = 1 – e^-0.883*coverage^*. Because of mis-mapping, read counts greater than zero do not necessarily imply that a MAG is present in the sample (Olm et al., 2021). To consider a MAG as present and minimise false positives, we required mapped reads to cover ≥ 50% of sites in a MAG (observed breadth ≥ 50%), or observed breadth ≥ 25% and the ratio of observed breadth:expected breadth ≥ 0.8. We alternatively tested a more permissible filter (observed breadth ≥ 5% breadth and observed breadth:expected breadth ≥ 0.3).

To calculate relative abundance from raw read counts, we first divided the number of reads mapped to a MAG by the total number of reads in the metagenomic sample, and then normalised this by the length of the MAG (to control for the varying lengths of MAGs). We then divided the resultant normalised read count of each MAG by the sum of normalised counts in the metagenomic sample to get the relative abundance. MAG prevalence was calculated by enumerating the number of samples a MAG was present in, and subsequently dividing by the total sample size. MAG social niche breadth (von Meijenfeldt et al., 2023) was calculated from the relative abundance table, with mean pairwise dissimilarity based on the inverse Spearman’s rank correlation of species (MAGs).

For parity, all databases were dereplicated at 95% ANI (i.e., at species-level representation) when comparing mapping rates across databases. To assess redundancy and overlap between UHGG (4,744 species-level MAGs), IndoMEE (1,304 species-level MAGs) and SPMP (646 species-level MAGs) databases, we combined and dereplicated these databases at 95% ANI to construct merged reference databases. This yielded 4,860 and 4,941 species-level MAGs for IndoMEE+UHGG and IndoMEE+UHGG+SPMP databases respectively — resulting in marginal (sub-additive) additions in species-level MAGs and suggesting considerable redundancy with UHGG.

### Profiling novelty

To identify novel species and subspecies among our assembled MAGs, we queried our MAGs against the largest available reference dataset (UHGG v2.0.1, full dataset: 289,232 MAGs) using magscreen (Almeida, 2021b). Magscreen first calculates Mash distances (Ondov et al., 2016) between query MAGs and reference MAGs (i.e., in the reference database) to identify the closest reference MAG (to the query MAG) and, based on this and the alignment fraction as calculated using MUMmer, calculates the average nucleotide identity (ANI) of the query MAG and its nearest reference MAG. Following dereplication (dRep; -pa 0.9 -sa 0.95 -nc 0.30 -cm larger -comp 50 -con 5) and GUNC based quality-control filtering (same parameters as detailed in Metagenomic assembly), a MAG is identified as novel at the species level if the ANI to the closest representative in the reference is less than 95%, and at the subspecies level if the corresponding ANI is less than 98%.

### Microbiome diversity and composition analysis

To investigate patterns in microbiome diversity across communities, we employed the microbiome relative abundance profiles of 116 Indonesian and 109 Singaporean individuals (SPMP), that were mapped on the IndoMEE species-level reference assembly (comprising 1,304 MAGs). To assess the effect of reference bias, we also calculated relative abundance profiles from mapping against the IndoMEE+UHGG+SPMP species-level merged assembly (comprising 4,941 MAGs). The Indonesian samples comprised members from i) remote communities: Asmat Daikot (Asmat_DAI, 5 samples), Punan Batu Sajau (Punan_PBS, 3 samples), Punan Tubu Hulu (Punan_HUL, 12 samples), and Punan Aput from Long Sule (Punan_SUL, 18 samples); ii) rural communities: Punan Tubu Respen (Punan_RPN, 20 samples), Basap (Basap_BRU, 31 samples), and Balinese in Pedawa (Balinese_PDW, 13 samples); and iii) an urban community: Balinese in Denpasar (Balinese_DPS, 14 samples). The urban Singaporean subset comprised samples from 3 ethnic groups: Chinese Singaporeans (Chinese_SIN, 53 samples), Malay Singaporeans (Malay_SIN, 26 samples), and Indian Singaporeans (Indian_SIN, 30 samples). Alpha diversity was estimated via three metrics: species richness, Shannon diversity index, and Faith’s phylogenetic diversity. To control for the effect of sequencing depth in our estimates of alpha diversity, we randomly downsampled all samples to a depth of 29.8 million reads per sample.

To assess differences in microbiome composition across samples, we calculated microbiome dissimilarity (beta diversity) using Aitchison distance (Aitchison et al., 2000), which applies a central-log transformation (CLR) on the relative abundance data. We did this separately for the Indonesia-Singapore dataset and the Indonesia-only dataset, and visualised the results using principal coordinate analysis (PCoA). Alpha and beta diversity were calculated from the relative abundance matrix using the abdiv v.0.2.0 (Bittinger, 2020) and vegan v.2.6-4 (Oksanen et al., 2022) packages, respectively, and performed in R (v.4.4.0).

### Microbiome cluster analysis

To identify the most appropriate number of clusters, we evaluated the stability and consistency of sample-to-cluster assignments across two datasets: the Indonesia-Singapore dataset (n = 225), and the Indonesia-only dataset (n = 116). Initial clustering was performed on Indonesia-Singapore using hierarchical clustering with the Ward.D2 linkage method. To define the upper limit of candidate cluster numbers, we applied the Elbow method to the within-cluster sum of squares (WSS) across k values from 1 to 15, identifying an inflection point at k = 4. Subsequently, the Gap Statistic (Tibshirani et al., 2001) was used to evaluate cluster separation and determine the optimal number of clusters within the range of k = 1 to 4. The same procedure was repeated for the Indonesia-only dataset. In both datasets, the Gap Statistic identified k = 2 as the optimal solution, though, given the different sample composition, these clusters cannot fully overlap.

Given that Indonesia is the central focus of this study, we sought to select a clustering configuration in the Indonesia-Singapore dataset that best preserved the k = 2 structure observed within the Indonesia-only dataset. To do this, we computed Adjusted Rand Index (ARI) values between each pair of cluster assignments over the two datasets. Among these, the k = 3 configuration in the Indonesia-Singapore dataset yielded the highest ARI value to the k = 2 solution in the Indonesia-only dataset (ARI = 0.519, compared to 0.03 for k = 1, 0.50 for k = 2, and 0.46 for k = 4), indicating a moderate level of concordance. Based on this integrative approach, we selected k = 3 as the optimal clustering solution for the full Indonesia-Singapore dataset.

To test for family-level differences in abundance between clusters, we performed Kruskal-Wallis tests with adjustments for multiple comparisons using the false discovery rate method. Additionally, to examine the divergence of urban Cluster 3 relative to the others (Cluster 1 and 2), we calculated the fold-change (FC) difference in abundance as 2^abs(log2(FC))^, where log_2_(FC) = log_2_(mean_Cluster3_ + 1 / mean_Cluster2+Cluster1_ + 1), and mean_Cluster_*i*_ is the mean family-level abundance in cluster *i*. Positive FC values indicate higher abundance in Cluster 3 while negative values indicate lower abundance.

### PERMANOVA model selection

To evaluate the contribution of predictor variables to microbiome composition (measured as Aitchison distance), we performed permutational multivariate analysis of variance (PERMANOVA) with model selection using the AICcPermanova package in R (Corcoran, 2023). The analysis was conducted on two datasets: the Indonesia-Singapore dataset (n = 225) and the Indonesia-only dataset (n = 116). We first assessed multicollinearity among these variables: lifestyle (ordinal variable; remote = 0, rural = 0.5, urban = 1), age, sex, population, study protocol (i.e. Singapore protocol, Bali protocol, and non-Bali Indonesian protocol), and the number of years since collection; by generating all possible models using the Aitchison distance matrix as the response. For each model, we calculated the Variance Inflation Factor (VIF); models with VIF values ≥ 6 were flagged as highly collinear and excluded from further analysis. Remaining models were then fitted for all combinations of predictors and ranked based on corrected Akaike Information Criterion (AICc), following the approach described previously (Burnham & Anderson, 2004). Across these models, we calculated AIC weights (i.e., the probability that a model is the best in terms of expected information loss among the candidate set) and used them for model averaging to estimate the adjusted R² of each predictor. The final output is a list of variables with their corresponding Akaike-adjusted R² values (‘NA’ if a variable has high collinearity, and weaker effect than that of its collinear variable(s)), representing their relative importance in explaining microbiome compositional variation while accounting for model complexity and sample size (Supplementary Table S4).

### Microbiome association with lifestyle cline

To investigate MAG association with the lifestyle cline, we conducted differential abundance analysis using MaAsLin2 (Mallick et al., 2021) under the model: MAG_abundance_ ∼ lifestyle_cline_ + age + sex; where lifestyle cline is defined as an ordinal variable (remote = 0, rural = 0.5, urban = 1) to reflect a linear gradient of lifestyle across communities (Supplementary Table S1), and age and sex are included as covariates. In this analysis, we required MAGs to be present in > 20 individuals (542 of the total 1,304 MAGs). We repeated this MaAsLin2 analysis for the Indonesia-only subset (n = 108; with prevalence filter removed to include all previously tested MAGs) as a robustness check.

### Microbiome association with population and lifestyle factors in Indonesia

To further assess the influence of lifestyle, we leveraged and analysed diet and activity data available for the Indonesian communities. Diet and activity-based distances between individuals were first visualised using principal component analysis (PCA) on 122 individuals with complete dietary data across five communities: Asmat Daikot, Punan Tubu Hulu, Basap, Balinese in Pedawa, and Balinese in Denpasar. To assess the broad association between lifestyle and microbiome composition, we performed a linear regression between the first principal component of the lifestyle PCA and the first principal coordinate of the microbiome PCoA, based on 73 individuals with complete metagenomic and lifestyle data. To evaluate population-specific associations, we additionally performed a linear mixed-model regression including population as a random effect, as well as separate linear regressions for each population.

To assess the correlation of microbiome distance with lifestyle distance within-and between-populations, we calculated Aitchison distances (based on MAGs with prevalence ≥ 2) and multivariate Euclidean distances of lifestyle factors (comprising the consumption of rice, sago, pork, chicken, fish, egg, and milk, along with participation in hunting and watching television) across pairs of individuals. Correlation strength and significance were subsequently evaluated using Mantel tests with 9,999 permutations.

To test for the effect of geography and lifestyle on microbiome divergence, we first calculated population mean relative abundances for MAGs and population mean lifestyle data. Pairwise distances were then calculated in terms of Aitchison distance for the former and 𝛘^2^ distance for the latter. Great-circle geographic distances between populations were measured using coordinates approximating the central location of each population. Total-permutation partial Mantel tests were then performed to assess the correlation between microbiome divergence and geographic distance accounting for lifestyle distance (i.e., *microbiome divergence ∼ geographic distance | lifestyle distance*), and between microbiome divergence and lifestyle distance accounting for geographic distance (*microbiome divergence ∼ lifestyle distance | geographic distance*).

To identify the factors driving MAG abundance, and disentangle their effects, we used MaAsLin2. We first ran MaAsLin2 with univariate models of diet (MAG_abundance_ ∼ d; where d = rice, sago, pork, egg, milk, fish or chicken) and population (MAG_abundance_ ∼ population) to quantify predictor importance, based on MAGs with prevalence > 5. This procedure identified 3 important diet variables: sago, pork and chicken. From this, we generated a more complete diet model (MAG_abundance_ ∼ sago + pork + chicken + age + sex + BMI) and compared this to a revised population model (MAG_abundance_ ∼ population + age + sex + BMI); with age, sex and BMI included in both models to control for potential confounding factors. We then compared the collective number of significant diet associations (diet-associated MAGs) to significant population associations (population-associated MAGs). This analysis captures to what extent diet or population is able to predict MAG relative abundance, and how often these signals overlap. Finally, to robustly identify true diet associations, we ran a mixed effects model incorporating the three dietary components as fixed effects and population as a random effect (MAG_abundance_ ∼ sago + pork + chicken + age + sex + BMI + 1|population), again controlling for potential confounding factors (age, sex and BMI). We calculated the variance explained by the population random effect (var_random_ = var_total_ - var_fixed_ - var_residual_) in this model to identify population-driven MAGs, and considered MAGs with a significant association to any of the three dietary factors as diet-driven. To account for potential methodological effects, we repeated this analysis for a subset of Indonesian samples (Asmat Daikot, Punan Tubu Hulu, and Basap, n = 47) that shared the same study protocol (Balinese samples were collected under a different protocol; see ‘Data and sample collection’ above), and obtained qualitatively consistent results.

### Functional annotation

To functionally annotate the IndoMEE reference database, we used genofan (Almeida, 2021a). Genofan first calls prokka version 1.14.6 (Seemann, 2014) to identify coding regions and annotate protein-coding genes, before annotating identified genes across a broad suite of functional classes via: AMRFinder version 3.11.14 for antimicrobial resistance (Feldgarden et al., 2021), gutSMASH version 1.0.0 for primary metabolic gene pathways (Andreu et al., 2023), antiSMASH version 6.0.1 for secondary metabolite biosynthetic gene clusters (Blin et al., 2021), dbCAN2 version 2.0.11 for carbohydrate-active enzyme and substrates (CAZy) (Zhang et al., 2018), KOFams (Aramaki et al., 2019) leveraging HMMER version 3.4 (Eddy, 2011) for Kyoto Encyclopedia of Genes and Genomes (KEGG) functional orthologs, and eggNOG-mapper version 2.1.3 (Huerta-Cepas et al., 2018) for gene ontologies (GO), clusters of orthologous genes (COG), protein families (Pfam), and others. Genofan and constituent tools were run under default settings.

### Functional enrichment analysis

To identify which functions or genes were significantly enriched or depleted in the novel MAGs relative to the known MAGs, we performed Fisher’s exact test on gutSMASH, antiSMASH, CAZy, AMRFinder, Pfam, GO and KEGG annotated features. These functional features were considered as present or absent and tabulated in 2 x 2 contingency tables of (novel, known):(present, absent), for each functional feature or gene. For COG, functional enrichment was assessed using Wilcoxon rank-sum test (two-sample, unpaired), given that COG categories are broader and return numerous hits per MAG; and hence better considered as counts data. For both sets of tests, we filtered out MAGs below 90% completeness, given that varying rates of completeness may bias enrichment results. For tests based on feature counts (i.e., COG), we normalised counts by MAG length (total feature counts), to control for varying MAG lengths. Test-inferred significance was adjusted for False Discovery Rate (FDR) and final significance was taken as that with FDR-adjusted *p* < 0.05.

### Functional composition and structure

To visualise compositional patterns in function across samples, we calculated KEGG, Pfam, CAZy, AMRFinder, gutSMASH, and antiSMASH (gene) counts for each MAG. Gene counts were weighted by the abundance of the MAG in the sample (individual). A centred log-ratio (CLR) transformation was applied to this table of abundance-weighted gene counts, to map compositional (i.e., Aitchison simplex space to Euclidean space), before applying an unscaled PCA. Mathematically, this is equivalent to applying a principal coordinate analysis on the Aitchison distance matrix.

## References

1. Abdill, R. J., Adamowicz, E. M., & Blekhman, R. (2022). Public human microbiome data are dominated by highly developed countries. PLOS Biology, 20(2), e3001536. 10.1371/journal.pbio.3001536

2. Aitchison, J., Barceló-Vidal, C., Martín-Fernández, J. A., & Pawlowsky-Glahn, V. (2000). Logratio Analysis and Compositional Distance. Mathematical Geology, 32(3), 271–275. 10.1023/A:1007529726302

3. Almeida, A. (2021a). *GenoFan—Genome functional annotation pipeline* [Computer software]. https://github.com/alexmsalmeida/genofan

4. Almeida, A. (2021b). *MAGscreen—Discovering new microbial species* [Computer software]. https://github.com/alexmsalmeida/magscreen

5. Almeida, A. (2021c). *metaMap—Quantifying genomes in metagenomes’* [Computer software]. https://github.com/alexmsalmeida/metamap

6. Almeida, A., Nayfach, S., Boland, M., Strozzi, F., Beracochea, M., Shi, Z. J., Pollard, K. S., Sakharova, E., Parks, D. H., Hugenholtz, P., Segata, N., Kyrpides, N. C., & Finn, R. D. (2021). A unified catalog of 204,938 reference genomes from the human gut microbiome. Nature Biotechnology, 39(1), 105–114. 10.1038/s41587-020-0603-3

7. Andreu, V. P., Augustijn, H. E., Chen, L., Zhernakova, A., Fu, J., Fischbach, M. A., Dodd, D., & Medema, M. H. (2023). gutSMASH predicts specialized primary metabolic pathways from the human gut microbiota. Nature Biotechnology, 41(10), 1416–1423. 10.1038/s41587-023-01675-1

8. Andreu-Sánchez, S., Blanco-Míguez, A., Wang, D., Golzato, D., Manghi, P., Heidrich, V., Fackelmann, G., Zhernakova, D. V., Kurilshikov, A., Valles-Colomer, M., Weersma, R. K., Zhernakova, A., Fu, J., & Segata, N. (2025). Global genetic diversity of human gut microbiome species is related to geographic location and host health. Cell, 188(15), 3942–3959.e9. 10.1016/j.cell.2025.04.014

9. Angelakis, E., Bachar, D., Yasir, M., Musso, D., Djossou, F., Gaborit, B., Brah, S., Diallo, A., Ndombe, G. M., Mediannikov, O., Robert, C., Azhar, E. I., Bibi, F., Nsana, N. S., Parra, H.-J., Akiana, J., Sokhna, C., Davoust, B., Dutour, A., & Raoult, D. (2019). Treponema species enrich the gut microbiota of traditional rural populations but are absent from urban individuals. New Microbes and New Infections, 27, 14–21. 10.1016/j.nmni.2018.10.009

10. Aramaki, T., Blanc-Mathieu, R., Endo, H., Ohkubo, K., Kanehisa, M., Goto, S., & Ogata, H. (2019). KofamKOALA: KEGG Ortholog assignment based on profile HMM and adaptive score threshold. Bioinformatics, 36(7), 2251–2252. 10.1093/bioinformatics/btz859

11. Badal, V. D., Vaccariello, E. D., Murray, E. R., Yu, K. E., Knight, R., Jeste, D. V., & Nguyen, T. T. (2020). The Gut Microbiome, Aging, and Longevity: A Systematic Review. Nutrients, 12(12), 3759. 10.3390/nu12123759

12. Bittinger, K. (2020). *abdiv: Alpha and beta diversity measures for community ecology* [Computer software]. https://cran.r-project.org/web/packages/abdiv/index.html.

13. Blin, K., Shaw, S., Kloosterman, A. M., Charlop-Powers, Z., van Wezel, G. P., Medema, M. H., & Weber, T. (2021). antiSMASH 6.0: Improving cluster detection and comparison capabilities. Nucleic Acids Research, 49(W1), W29–W35. 10.1093/nar/gkab335

14. Brito, I. L., Gurry, T., Zhao, S., Huang, K., Young, S. K., Shea, T. P., Naisilisili, W., Jenkins, A. P., Jupiter, S. D., Gevers, D., & Alm, E. J. (2019). Transmission of human-associated microbiota along family and social networks. Nature Microbiology, 4(6), Article 6. 10.1038/s41564-019-0409-6

15. Browne, H. P., Almeida, A., Kumar, N., Vervier, K., Adoum, A. T., Viciani, E., Dawson, N. J. R., Forster, S. C., Cormie, C., Goulding, D., & Lawley, T. D. (2021). Host adaptation in gut Firmicutes is associated with sporulation loss and altered transmission cycle. Genome Biology, 22(1), 204. 10.1186/s13059-021-02428-6

16. Burnham, K. P., & Anderson, D. R. (Eds.). (2004). Model Selection and Multimodel Inference. Springer. 10.1007/b97636

17. Cade, J. E., Burley, V. J., Warm, D. L., Thompson, R. L., & Margetts, B. M. (2004). Food-frequency questionnaires: A review of their design, validation and utilisation. Nutrition Research Reviews, 17(1), 5–22. 10.1079/NRR200370

18. Carter, M. M., Olm, M. R., Merrill, B. D., Dahan, D., Tripathi, S., Spencer, S. P., Yu, F. B., Jain, S., Neff, N., Jha, A. R., Sonnenburg, E. D., & Sonnenburg, J. L. (2023). Ultra-deep sequencing of Hadza hunter-gatherers recovers vanishing gut microbes. Cell, 186(14), 3111–3124.e13. 10.1016/j.cell.2023.05.046

19. Chklovski, A., Parks, D. H., Woodcroft, B. J., & Tyson, G. W. (2023). CheckM2: A rapid, scalable and accurate tool for assessing microbial genome quality using machine learning. Nature Methods, 20(8), 1203–1212. 10.1038/s41592-023-01940-w

20. Chonnacháin, C. N., Feeney, E. L., Gollogly, C., Shields, D. C., Loscher, C. E., Cotter, P. D., Noronha, N., Stack, R., Doherty, G. A., & Gibney, E. R. (2024). The effects of dairy on the gut microbiome and symptoms in gastrointestinal disease cohorts: A systematic review. Gut Microbiome, 5, e5. 10.1017/gmb.2024.2

21. Corcoran, D. (2023). *AICcPermanova: Model Selection of PERMANOVA Models Using AICc* [Computer software]. 10.5281/zenodo.7816863

22. Danecek, P., Bonfield, J. K., Liddle, J., Marshall, J., Ohan, V., Pollard, M. O., Whitwham, A., Keane, T., McCarthy, S. A., Davies, R. M., & Li, H. (2021). Twelve years of SAMtools and BCFtools. GigaScience, 10(2), giab008. 10.1093/gigascience/giab008

23. David, L. A., Maurice, C. F., Carmody, R. N., Gootenberg, D. B., Button, J. E., Wolfe, B. E., Ling, A. V., Devlin, A. S., Varma, Y., Fischbach, M. A., Biddinger, S. B., Dutton, R. J., & Turnbaugh, P. J. (2014). Diet rapidly and reproducibly alters the human gut microbiome. Nature, 505(7484), Article 7484. 10.1038/nature12820

24. Debray, R., Tung, J., & Archie, E. A. (2024). Ecology and Evolution of the Social Microbiome. Annual Review of Ecology, Evolution, and Systematics, 55(Volume 55, 2024), 89–114. 10.1146/annurev-ecolsys-102622-030749

25. Delgado, L. F., & Andersson, A. F. (2022). Evaluating metagenomic assembly approaches for biome-specific gene catalogues. Microbiome, 10(1), 72. 10.1186/s40168-022-01259-2

26. Derrien, M., Turroni, F., Ventura, M., & van Sinderen, D. (2022). Insights into endogenous Bifidobacterium species in the human gut microbiota during adulthood. Trends in Microbiology, 30(10), 940–947. 10.1016/j.tim.2022.04.004

27. Dounias, E., & Froment, A. (2011). From foraging to farming among present-day forest hunter-gatherers: Consequences on diet and health. International Forestry Review, 13(3), 294–304. 10.1505/146554811798293818

28. Dounias, E., Selzner, A., Koizumi, M., & Levang, P. (2007). From Sago to Rice, from Forest to Town: The Consequences of Sedentarization for the Nutritional Ecology of Punan Former Hunter-Gatherers of Borneo. Food and Nutrition Bulletin, 28(2_suppl2), S294–S302. 10.1177/15648265070282S208

29. Eddy, S. R. (2011). Accelerated Profile HMM Searches. PLOS Computational Biology, 7(10), e1002195. 10.1371/journal.pcbi.1002195

30. Febinia, C. A., Malik, S. G., Djuwita, R., Weta, I. W., Wihandani, D. M., Maulida, R., Sudoyo, H., & Holmes, A. J. (2022). Distinctive Microbiome Type Distribution in a Young Adult Balinese Cohort May Reflect Environmental Changes Associated with Modernization. Microbial Ecology, 83(3), 798–810. 10.1007/s00248-021-01786-9

31. Feldgarden, M., Brover, V., Gonzalez-Escalona, N., Frye, J. G., Haendiges, J., Haft, D. H., Hoffmann, M., Pettengill, J. B., Prasad, A. B., Tillman, G. E., Tyson, G. H., & Klimke, W. (2021). AMRFinderPlus and the Reference Gene Catalog facilitate examination of the genomic links among antimicrobial resistance, stress response, and virulence. Scientific Reports, 11(1), 12728. 10.1038/s41598-021-91456-0

32. Galperin, M. Y. (2013). Genome Diversity of Spore-Forming Firmicutes. Microbiology Spectrum, 1(2), 10.1128/microbiolspectrum.tbs-0015–2012.10.1128/microbiolspectrum.tbs-0015-2012

33. Gomez, A., Petrzelkova, K. J., Burns, M. B., Yeoman, C. J., Amato, K. R., Vlckova, K., Modry, D., Todd, A., Jost Robinson, C. A., Remis, M. J., Torralba, M. G., Morton, E., Umaña, J. D., Carbonero, F., Gaskins, H. R., Nelson, K. E., Wilson, B. A., Stumpf, R. M., White, B. A., … Blekhman, R. (2016). Gut Microbiome of Coexisting BaAka Pygmies and Bantu Reflects Gradients of Traditional Subsistence Patterns. Cell Reports, 14(9), 2142–2153. 10.1016/j.celrep.2016.02.013

34. Gorvitovskaia, A., Holmes, S. P., & Huse, S. M. (2016). Interpreting Prevotella and Bacteroides as biomarkers of diet and lifestyle. Microbiome, 4, 15. 10.1186/s40168-016-0160-7

35. Gounot, J.-S., Chia, M., Bertrand, D., Saw, W.-Y., Ravikrishnan, A., Low, A., Ding, Y., Ng, A. H. Q., Tan, L. W. L., Teo, Y.-Y., Seedorf, H., & Nagarajan, N. (2022). Genome-centric analysis of short and long read metagenomes reveals uncharacterized microbiome diversity in Southeast Asians. Nature Communications, 13(1), Article 1. 10.1038/s41467-022-33782-z

36. Guan, H., Pu, Y., Liu, C., Lou, T., Tan, S., Kong, M., Sun, Z., Mei, Z., Qi, Q., Quan, Z., Zhao, G., & Zheng, Y. (2021). Comparison of fecal collection methods on variation in gut metagenomics and untargeted metabolomics. mSphere, 6(1), 00636–21. 10.1128/msphere.00636-21

37. Gupta, V. K., Paul, S., & Dutta, C. (2017). Geography, Ethnicity or Subsistence-Specific Variations in Human Microbiome Composition and Diversity. Frontiers in Microbiology, 8, 1162. 10.3389/fmicb.2017.01162

38. Huerta-Cepas, J., Szklarczyk, D., Heller, D., Hernández-Plaza, A., Forslund, S. K., Cook, H., Mende, D. R., Letunic, I., Rattei, T., Jensen, L. J., Mering, C., & Bork, P. (2018). eggNOG 5.0: A hierarchical, functionally and phylogenetically annotated orthology resource based on 5090 organisms and 2502 viruses. Nucleic Acids Research, 47(D1), 309–314. 10.1093/nar/gky1085

39. Ilett, E. E., Jørgensen, M., & Noguera-Julian, M. (2019). Gut microbiome comparability of fresh-frozen versus stabilized-frozen samples from hospitalized patients using 16S rRNA gene and shotgun metagenomic sequencing. Scientific Reports, 9, 13351. 10.1038/s41598-019-49956-7

40. Jain, C., Rodriguez-R, L. M., Phillippy, A. M., Konstantinidis, K. T., & Aluru, S. (2018). High throughput ANI analysis of 90K prokaryotic genomes reveals clear species boundaries. Nature Communications, 9(1), 5114. 10.1038/s41467-018-07641-9

41. Jha, A. R., Davenport, E. R., Gautam, Y., Bhandari, D., Tandukar, S., Ng, K. M., Fragiadakis, G. K., Holmes, S., Gautam, G. P., Leach, J., Sherchand, J. B., Bustamante, C. D., & Sonnenburg, J. L. (2018). Gut microbiome transition across a lifestyle gradient in Himalaya. PLOS Biology, 16(11), e2005396. 10.1371/journal.pbio.2005396

42. Johnson, A. J., Zheng, J. J., Kang, J. W., Saboe, A., Knights, D., & Zivkovic, A. M. (2020). A Guide to Diet-Microbiome Study Design. Frontiers in Nutrition, 7. 10.3389/fnut.2020.00079

43. Jung, Y., & Han, D. (2022). BWA-MEME: BWA-MEM emulated with a machine learning approach. Bioinformatics, 38(9), 2404–2413. 10.1093/bioinformatics/btac137

44. Korpela, K., Salonen, A., Virta, L. J., Kekkonen, R. A., Forslund, K., Bork, P., & de Vos, W. M. (2016). Intestinal microbiome is related to lifetime antibiotic use in Finnish pre-school children. Nature Communications, 7(1), 10410. 10.1038/ncomms10410

45. Krueger, F. (2016). *Babraham Bioinformatics—Trim Galore* [Computer software]. https://github.com/FelixKrueger/TrimGalore

46. Leviatan, S., Shoer, S., Rothschild, D., Gorodetski, M., & Segal, E. (2022). An expanded reference map of the human gut microbiome reveals hundreds of previously unknown species. Nature Communications, 13(1), 3863. 10.1038/s41467-022-31502-1

47. Li, H. (2013). *Aligning sequence reads, clone sequences and assembly contigs with BWA-MEM* (arXiv:1303.3997). arXiv. 10.48550/arXiv.1303.3997

48. Lu, J., Zhang, L., Zhang, H., Chen, Y., Zhao, J., Chen, W., Lu, W., & Li, M. (2023). Population-level variation in gut bifidobacterial composition and association with geography, age, ethnicity, and staple food. Npj Biofilms and Microbiomes, 9(1), 1–12. 10.1038/s41522-023-00467-4

49. Mallick, H., Rahnavard, A., McIver, L. J., Ma, S., Zhang, Y., Nguyen, L. H., Tickle, T. L., Weingart, G., Ren, B., Schwager, E. H., Chatterjee, S., Thompson, K. N., Wilkinson, J. E., Subramanian, A., Lu, Y., Waldron, L., Paulson, J. N., Franzosa, E. A., Bravo, H. C., & Huttenhower, C. (2021). Multivariable association discovery in population-scale meta-omics studies. PLOS Computational Biology, 17(11), e1009442. 10.1371/journal.pcbi.1009442

50. Mancabelli, L., Milani, C., Lugli, G. A., Turroni, F., Ferrario, C., van Sinderen, D., & Ventura, M. (2017). Meta-analysis of the human gut microbiome from urbanized and pre-agricultural populations. Environmental Microbiology, 19(4), 1379–1390. 10.1111/1462-2920.13692

51. Nagata, N., Nishijima, S., Miyoshi-Akiyama, T., Kojima, Y., Kimura, M., Aoki, R., Ohsugi, M., Ueki, K., Miki, K., Iwata, E., Hayakawa, K., Ohmagari, N., Oka, S., Mizokami, M., Itoi, T., Kawai, T., Uemura, N., & Hattori, M. (2022). Population-level Metagenomics Uncovers Distinct Effects of Multiple Medications on the Human Gut Microbiome. Gastroenterology, 163(4), 1038–1052. 10.1053/j.gastro.2022.06.070

52. Na’im, A., & Syaputra, H. (2012). Kewarganegaraan Suku Bangsa Agama dan Bahasa Sehari-hari Penduduk Indonesia [Nationality, Ethnicity, Religion, and Daily Language of Indonesian Population]. Badan Pusat Statistik [BPS-Statistics Indonesia]. https://www.bps.go.id/en/publication/2012/05/23/55eca38b7fe0830834605b35/kewarganegaraan-suku-bangsa-agama-dan-bahasa-sehari-hari-penduduk-indonesia.html

53. Nakayama, J., Watanabe, K., Jiang, J., Matsuda, K., Chao, S.-H., Haryono, P., La-ongkham, O., Sarwoko, M.-A., Sujaya, I. N., Zhao, L., Chen, K.-T., Chen, Y.-P., Chiu, H.-H., Hidaka, T., Huang, N.-X., Kiyohara, C., Kurakawa, T., Sakamoto, N., Sonomoto, K., … Lee, Y.-K. (2015). Diversity in gut bacterial community of school-age children in Asia. Scientific Reports, 5(1), 8397. 10.1038/srep08397

54. Nguyen, L.-T., Schmidt, H. A., von Haeseler, A., & Minh, B. Q. (2015). IQ-TREE: A Fast and Effective Stochastic Algorithm for Estimating Maximum-Likelihood Phylogenies. Molecular Biology and Evolution, 32(1), 268–274. 10.1093/molbev/msu300

55. Obregon-Tito, A. J., Tito, R. Y., Metcalf, J., Sankaranarayanan, K., Clemente, J. C., Ursell, L. K., Zech Xu, Z., Van Treuren, W., Knight, R., Gaffney, P. M., Spicer, P., Lawson, P., Marin-Reyes, L., Trujillo-Villarroel, O., Foster, M., Guija-Poma, E., Troncoso-Corzo, L., Warinner, C., Ozga, A. T., & Lewis, C. M. (2015). Subsistence strategies in traditional societies distinguish gut microbiomes. Nature Communications, 6(1), Article 1. 10.1038/ncomms7505

56. Oksanen, J., Simpson, G. L., Blanchet, F. G., Kindt, R., Legendre, P., Minchin, P. R., O’Hara, R. B., Solymos, P., Stevens, M. H. H., Szoecs, E., Wagner, H., Barbour, M., Bedward, M., Bolker, B., Borcard, D., Carvalho, G., Chirico, M., Caceres, M. D., Durand, S., … Weedon, J. (2022). *vegan: Community Ecology Package.* (Version R package version 2.7-1) [Computer software]. https://CRAN.R-project.org/package=vegan

57. Olm, M. R., Brown, C. T., Brooks, B., & Banfield, J. F. (2017). dRep: A tool for fast and accurate genomic comparisons that enables improved genome recovery from metagenomes through de-replication. The ISME Journal, 11(12), 2864–2868. 10.1038/ismej.2017.126

58. Olm, M. R., Crits-Christoph, A., Bouma-Gregson, K., Firek, B. A., Morowitz, M. J., & Banfield, J. F. (2021). inStrain profiles population microdiversity from metagenomic data and sensitively detects shared microbial strains. Nature Biotechnology, 39(6), 727–736. 10.1038/s41587-020-00797-0

59. Ondov, B. D., Treangen, T. J., Melsted, P., Mallonee, A. B., Bergman, N. H., Koren, S., & Phillippy, A. M. (2016). Mash: Fast genome and metagenome distance estimation using MinHash. Genome Biology, 17(1), 132. 10.1186/s13059-016-0997-x

60. Orakov, A., Fullam, A., Coelho, L. P., Khedkar, S., Szklarczyk, D., Mende, D. R., Schmidt, T. S. B., & Bork, P. (2021). GUNC: Detection of chimerism and contamination in prokaryotic genomes. Genome Biology, 22(1), 178. 10.1186/s13059-021-02393-0

61. Parks, D. H., Imelfort, M., Skennerton, C. T., Hugenholtz, P., & Tyson, G. W. (2015). CheckM: Assessing the quality of microbial genomes recovered from isolates, single cells, and metagenomes. Genome Research, 25(7), 1043–1055. 10.1101/gr.186072.114

62. Pasolli, E., Asnicar, F., Manara, S., Zolfo, M., Karcher, N., Armanini, F., Beghini, F., Manghi, P., Tett, A., Ghensi, P., Collado, M. C., Rice, B. L., DuLong, C., Morgan, X. C., Golden, C. D., Quince, C., Huttenhower, C., & Segata, N. (2019). Extensive Unexplored Human Microbiome Diversity Revealed by Over 150,000 Genomes from Metagenomes Spanning Age, Geography, and Lifestyle. Cell, 176(3), 649–662.e20. 10.1016/j.cell.2019.01.001

63. Penders, J., Thijs, C., Vink, C., Stelma, F. F., Snijders, B., Kummeling, I., van den Brandt, P. A., & Stobberingh, E. E. (2006). Factors Influencing the Composition of the Intestinal Microbiota in Early Infancy. Pediatrics, 118(2), 511–521. 10.1542/peds.2005-2824

64. Popkin, B. M. (2004). The Nutrition Transition: An Overview of World Patterns of Change. Nutrition Reviews, 62(suppl_2), S140–S143. 10.1111/j.1753-4887.2004.tb00084.x

65. Popkin, B. M. (2011). Contemporary nutritional transition: Determinants of diet and its impact on body composition. Proceedings of the Nutrition Society, 70(1), 82–91. 10.1017/S0029665110003903

66. Rampelli, S., Turroni, S., & Candela, M. (2023). Ageing and Human Gut Microbiome: The Taxonomic and Functional Transition Towards an Elderly-Type Microbiome. In F. Marotta (Ed.), Gut Microbiota in Aging and Chronic Diseases (pp. 23–39). Springer International Publishing. 10.1007/978-3-031-14023-5_2

67. Reyes-García, V., Powell, B., Díaz-Reviriego, I., Fernández-Llamazares, Á., Gallois, S., & Gueze, M. (2019). Dietary transitions among three contemporary hunter-gatherers across the tropics. Food Security, 11(1), 109–122. 10.1007/s12571-018-0882-4

68. Rosas-Plaza, S., Hernández-Terán, A., Navarro-Díaz, M., Escalante, A. E., Morales-Espinosa, R., & Cerritos, R. (2022). Human Gut Microbiome Across Different Lifestyles: From Hunter-Gatherers to Urban Populations. Frontiers in Microbiology, 13. 10.3389/fmicb.2022.843170

69. Rothschild, D., Weissbrod, O., Barkan, E., Kurilshikov, A., Korem, T., Zeevi, D., Costea, P. I., Godneva, A., Kalka, I. N., Bar, N., Shilo, S., Lador, D., Vila, A. V., Zmora, N., Pevsner-Fischer, M., Israeli, D., Kosower, N., Malka, G., Wolf, B. C., … Segal, E. (2018). Environment dominates over host genetics in shaping human gut microbiota. Nature, 555(7695), 210–215. 10.1038/nature25973

70. Sarkar, A., Harty, S., Johnson, K. V.-A., Moeller, A. H., Archie, E. A., Schell, L. D., Carmody, R. N., Clutton-Brock, T. H., Dunbar, R. I. M., & Burnet, P. W. J. (2020). Microbial transmission in animal social networks and the social microbiome. Nature Ecology & Evolution, 4(8), 1020–1035. 10.1038/s41559-020-1220-8

71. Sarkar, A., McInroy, C. J. A., Harty, S., Raulo, A., Ibata, N. G. O., Valles-Colomer, M., Johnson, K. V.-A., Brito, I. L., Henrich, J., Archie, E. A., Barreiro, L. B., Gazzaniga, F. S., Finlay, B. B., Koonin, E. V., Carmody, R. N., & Moeller, A. H. (2024). Microbial transmission in the social microbiome and host health and disease. Cell, 187(1), 17–43. 10.1016/j.cell.2023.12.014

72. Schnorr, S. L., Candela, M., Rampelli, S., Centanni, M., Consolandi, C., Basaglia, G., Turroni, S., Biagi, E., Peano, C., Severgnini, M., Fiori, J., Gotti, R., De Bellis, G., Luiselli, D., Brigidi, P., Mabulla, A., Marlowe, F., Henry, A. G., & Crittenden, A. N. (2014). Gut microbiome of the Hadza hunter-gatherers. Nature Communications, 5(1), Article 1. 10.1038/ncomms4654

73. Seemann, T. (2014). Prokka: Rapid prokaryotic genome annotation. Bioinformatics, 30(14), 2068–2069. 10.1093/bioinformatics/btu153

74. Song, S. J., Amir, A., Metcalf, J. L., Amato, K. R., Xu, Z. Z., Humphrey, G., & Knight, R. (2016). Preservation methods differ in fecal microbiome stability, affecting suitability for field studies. mSystems, 1(3), 00021–16. 10.1128/msystems.00021-16.

75. Sonnenburg, E. D., & Sonnenburg, J. L. (2019). The ancestral and industrialized gut microbiota and implications for human health. Nature Reviews Microbiology, 17(6), Article 6. 10.1038/s41579-019-0191-8

76. Stražar, M., Temba, G. S., Vlamakis, H., Kullaya, V. I., Lyamuya, F., Mmbaga, B. T., Joosten, L. A. B., van der Ven, A. J. A. M., Netea, M. G., de Mast, Q., & Xavier, R. J. (2021). Gut microbiome-mediated metabolism effects on immunity in rural and urban African populations. Nature Communications, 12(1), 4845. 10.1038/s41467-021-25213-2

77. Suzuki, T. A., Fitzstevens, J. L., Schmidt, V. T., Enav, H., Huus, K. E., Mbong Ngwese, M., Grießhammer, A., Pfleiderer, A., Adegbite, B. R., Zinsou, J. F., Esen, M., Velavan, T. P., Adegnika, A. A., Song, L. H., Spector, T. D., Muehlbauer, A. L., Marchi, N., Kang, H., Maier, L., … Ley, R. E. (2022). Codiversification of gut microbiota with humans. Science, 377(6612), 1328–1332. 10.1126/science.abm7759

78. Swick, M. C., Koehler, T. M., & Driks, A. (2016). Surviving Between Hosts: Sporulation and Transmission. In Virulence Mechanisms of Bacterial Pathogens (pp. 567–591). John Wiley & Sons, Ltd. 10.1128/9781555819286.ch20

79. Tibshirani, R., Walther, G., & Hastie, T. (2001). Estimating the number of clusters in a data set via the gap statistic. Journal of the Royal Statistical Society: Series B (Statistical Methodology, 63(2), 411–423. 10.1111/1467-9868.00293

80. Uritskiy, G. V., DiRuggiero, J., & Taylor, J. (2018). MetaWRAP—a flexible pipeline for genome-resolved metagenomic data analysis. Microbiome, 6(1), 158. 10.1186/s40168-018-0541-1

81. Valdes, A. M., Walter, J., Segal, E., & Spector, T. D. (2018). Role of the gut microbiota in nutrition and health. BMJ, 361, k2179. 10.1136/bmj.k2179

82. Valles-Colomer, M., Blanco-Míguez, A., Manghi, P., Asnicar, F., Dubois, L., Golzato, D., Armanini, F., Cumbo, F., Huang, K. D., Manara, S., Masetti, G., Pinto, F., Piperni, E., Punčochář, M., Ricci, L., Zolfo, M., Farrant, O., Goncalves, A., Selma-Royo, M., … Segata, N. (2023). The person-to-person transmission landscape of the gut and oral microbiomes. Nature, 614(7946), 125–135. 10.1038/s41586-022-05620-1

83. Vangay, P., Johnson, A. J., Ward, T. L., Al-Ghalith, G. A., Shields-Cutler, R. R., Hillmann, B. M., Lucas, S. K., Beura, L. K., Thompson, E. A., Till, L. M., Batres, R., Paw, B., Pergament, S. L., Saenyakul, P., Xiong, M., Kim, A. D., Kim, G., Masopust, D., Martens, E. C., … Knights, D. (2018). US Immigration Westernizes the Human Gut Microbiome. Cell, 175(4), 962–972.e10. 10.1016/j.cell.2018.10.029

84. von Meijenfeldt, F. A. B., Hogeweg, P., & Dutilh, B. E. (2023). A social niche breadth score reveals niche range strategies of generalists and specialists. Nature Ecology & Evolution, 7(5), 768–781. 10.1038/s41559-023-02027-7

85. Wilson, A. S., Koller, K. R., Ramaboli, M. C., Nesengani, L. T., Ocvirk, S., Chen, C., Flanagan, C. A., Sapp, F. R., Merritt, Z. T., Bhatti, F., Thomas, T. K., & O’Keefe, S. J. D. (2020). Diet and the Human Gut Microbiome: An International Review. Digestive Diseases and Sciences, 65(3), 723–740. 10.1007/s10620-020-06112-w

86. Yatsunenko, T., Rey, F. E., Manary, M. J., Trehan, I., Dominguez-Bello, M. G., Contreras, M., Magris, M., Hidalgo, G., Baldassano, R. N., Anokhin, A. P., Heath, A. C., Warner, B., Reeder, J., Kuczynski, J., Caporaso, J. G., Lozupone, C. A., Lauber, C., Clemente, J. C., Knights, D., … Gordon, J. I. (2012). Human gut microbiome viewed across age and geography. Nature, 486(7402), Article 7402. 10.1038/nature11053

87. Zahorka, H. (2017). Punan ‘Gita,’ Penan benalui, Punan aput: From hunter-gatherers to average citizens: Early and later experiences of the author in East Kalimantan. Borneo Research Bulletin, 48, 283+. Gale Academic OneFile.

88. Zhang, H., Yohe, T., Huang, L., Entwistle, S., Wu, P., Yang, Z., Busk, P. K., Xu, Y., & Yin, Y. (2018). dbCAN2: A meta server for automated carbohydrate-active enzyme annotation. Nucleic Acids Research, 46(W1), W95–W101. 10.1093/nar/gky418

89. Zhernakova, A., Kurilshikov, A., Bonder, M. J., Tigchelaar, E. F., Schirmer, M., Vatanen, T., Mujagic, Z., Vila, A. V., Falony, G., Vieira-Silva, S., Wang, J., Imhann, F., Brandsma, E., Jankipersadsing, S. A., Joossens, M., Cenit, M. C., Deelen, P., Swertz, M. A., LifeLines cohort study, … Fu, J. (2016). Population-based metagenomics analysis reveals markers for gut microbiome composition and diversity. Science, 352(6285), 565–569. 10.1126/science.aad3369

