## Supplementary Document S1 for "From sporulation to village differentiation: the shaping of the social microbiome over rural-to-urban lifestyle transition in Indonesia"

### Supplementary Information

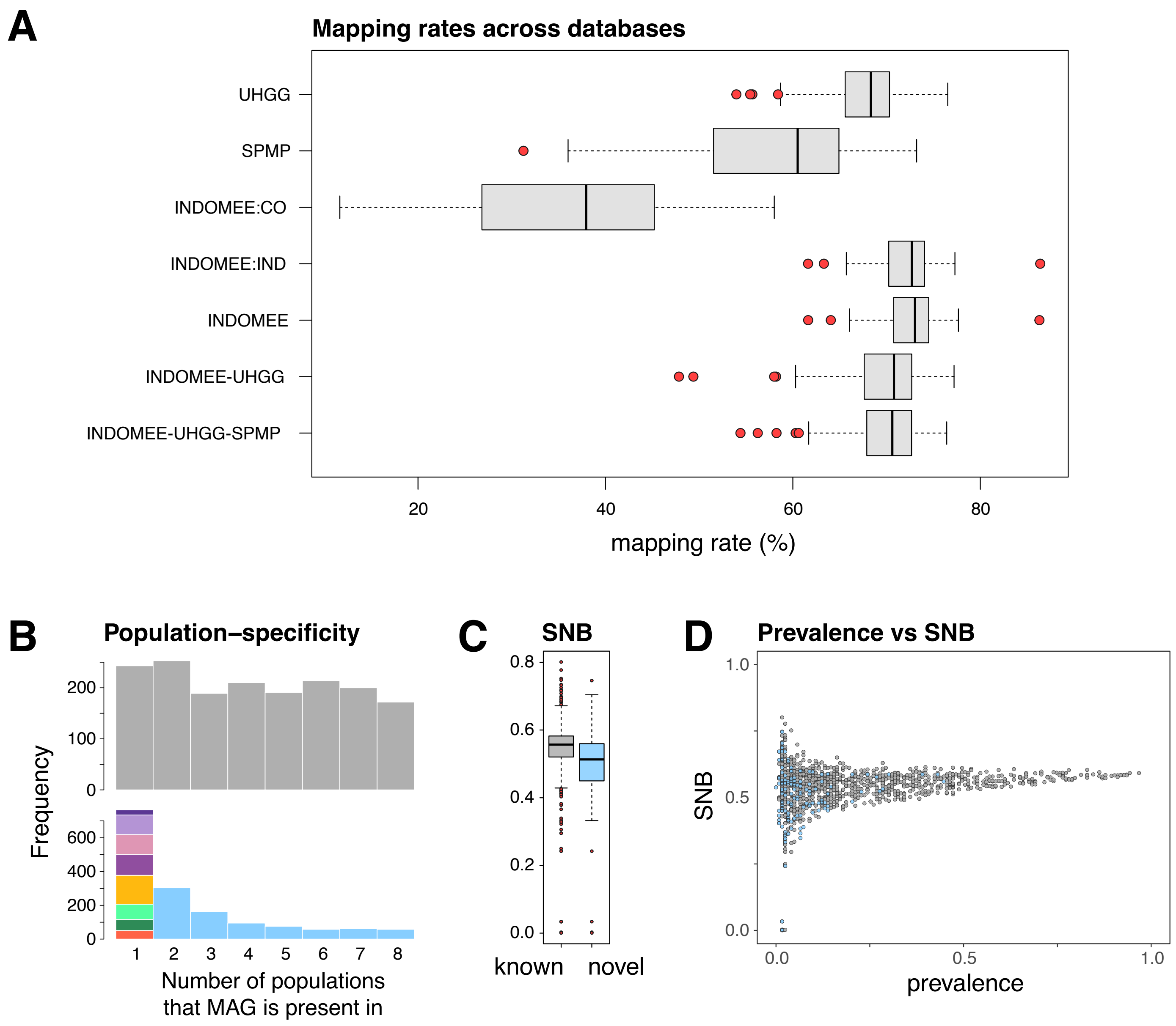


**Supplementary Figure S1. Characterisation of novelty, prevalence and database mapping rates.** **A**) Mapping rate of host-decontaminated metagenomic reads of 116 Indonesian samples across 7 genome-reference databases, dereplicated at 95% ANI. UHGG = Unified Human Gastrointestinal Genome (v2.0.1), SPMP = Singapore Platinum Metagenomes Project, INDOMEE:CO = IndoMEE co-assembled MAGs, INDOMEE:IND = IndoMEE individual-assembled MAGs, INDOMEE = IndoMEE individual- & co-assembled MAGs. Databases with hyphenated names indicate combined databases. **B**) Histograms of village-level prevalence for novel (bottom; coloured) and known (top; grey) subspecies-level MAGs. Novel MAGs (bottom panel; coloured) are, relatively, more often unique to a single village than known MAGs (top panel; grey) in the IndoMEE database (two-sample Kolmogorov-Smirnov test, D = 0.38, $p < 0.0001$). Colours for village-specific novel MAGs correspond to that in Figure 1. **C**) Social niche breadth (SNB) of novel MAGs compared to known MAGs, shown for species-level MAGs. Novel MAGs are found in more similar microbial compositions (i.e., exhibit a lower social niche breadth score) than known MAGs (Mann-Whitney two-sided test, $p < 0.0001$). **D**) Prevalence and SNB scores are plotted for novel (blue) and known (grey) species-level MAGs. For boxplots in **A**) and **C**), centre lines denote medians, box edges indicate lower and upper quartiles, and whiskers extend to 1.5 times the interquartile range.

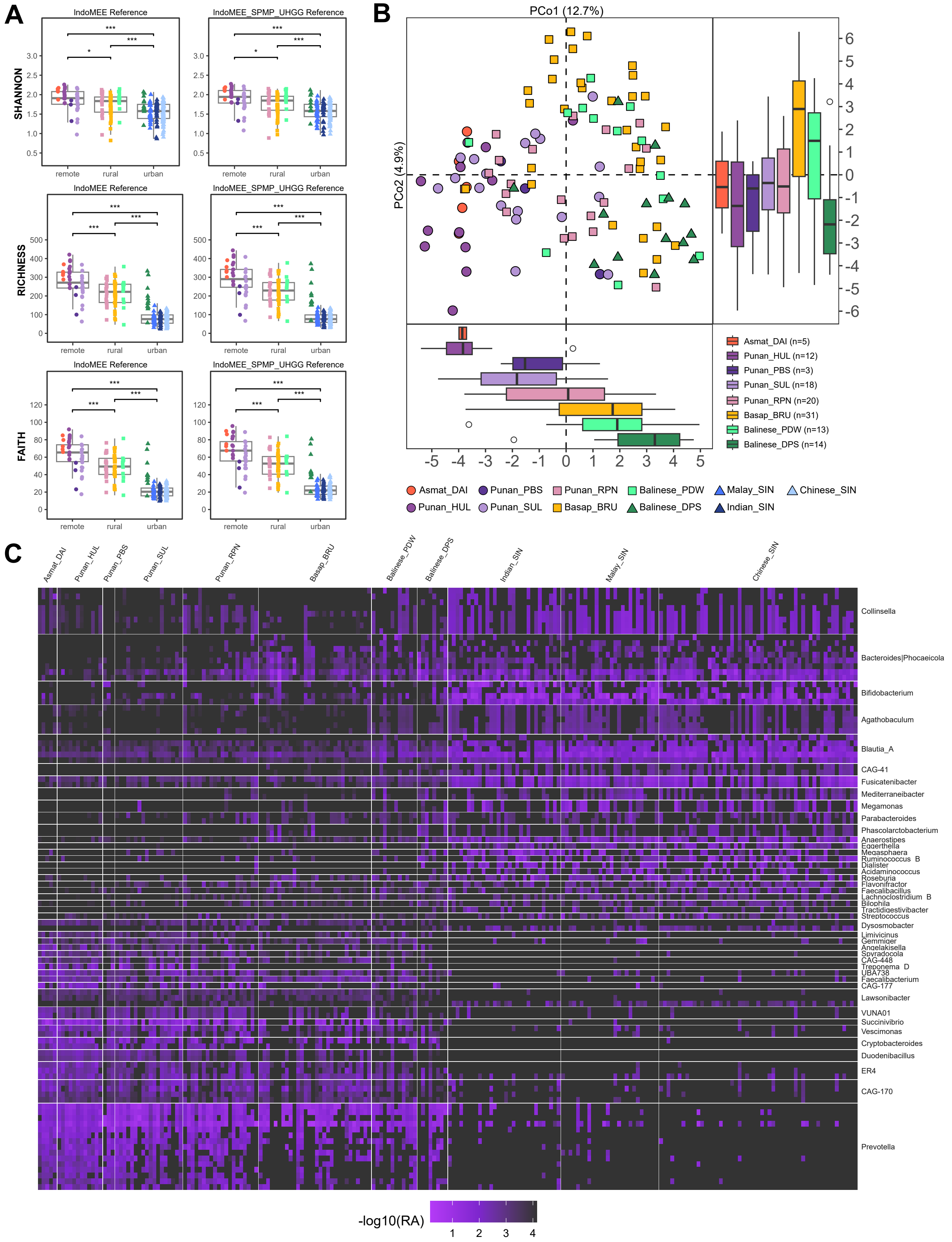


**Supplementary Figure S2. Microbiome variation in 116 Indonesian and 109 Singaporean samples across remote, rural, and urban populations. A)** Distribution of alpha diversity metrics (Shannon diversity index, species richness, and Faith's phylogenetic diversity) across lifestyle groups (remote, rural, urban). *, ** and *** denote significance thresholds of $p < 0.05$, $p < 0.01$, and $p < 0.001$, respectively. Centre lines denote medians, box edges indicate lower and upper quartiles, and whiskers extend to 1.5 times the interquartile range. **B)** Principal coordinate analysis of Indonesian faecal microbiome based on Aitchison distance of 1,079 MAGs present in >2 individuals. Box and whiskers plots show the positional distribution of individuals by population along the first and second principal coordinates. In box plots, centre lines denote medians, box edges indicate lower and upper quartiles, and whiskers extend to 1.5 times the interquartile range. **C)** Relative abundance heatmap (-$\log_{10}$ transformed) of the top 100 species most strongly associated with the lifestyle cline, depicting trends across and within genera. Taxa are ordered by their lifestyle cline (ordinal scoring: remote - 0, rural - 0.5, urban - 1) coefficients. See Supplementary Table S4 for additional species-level data.

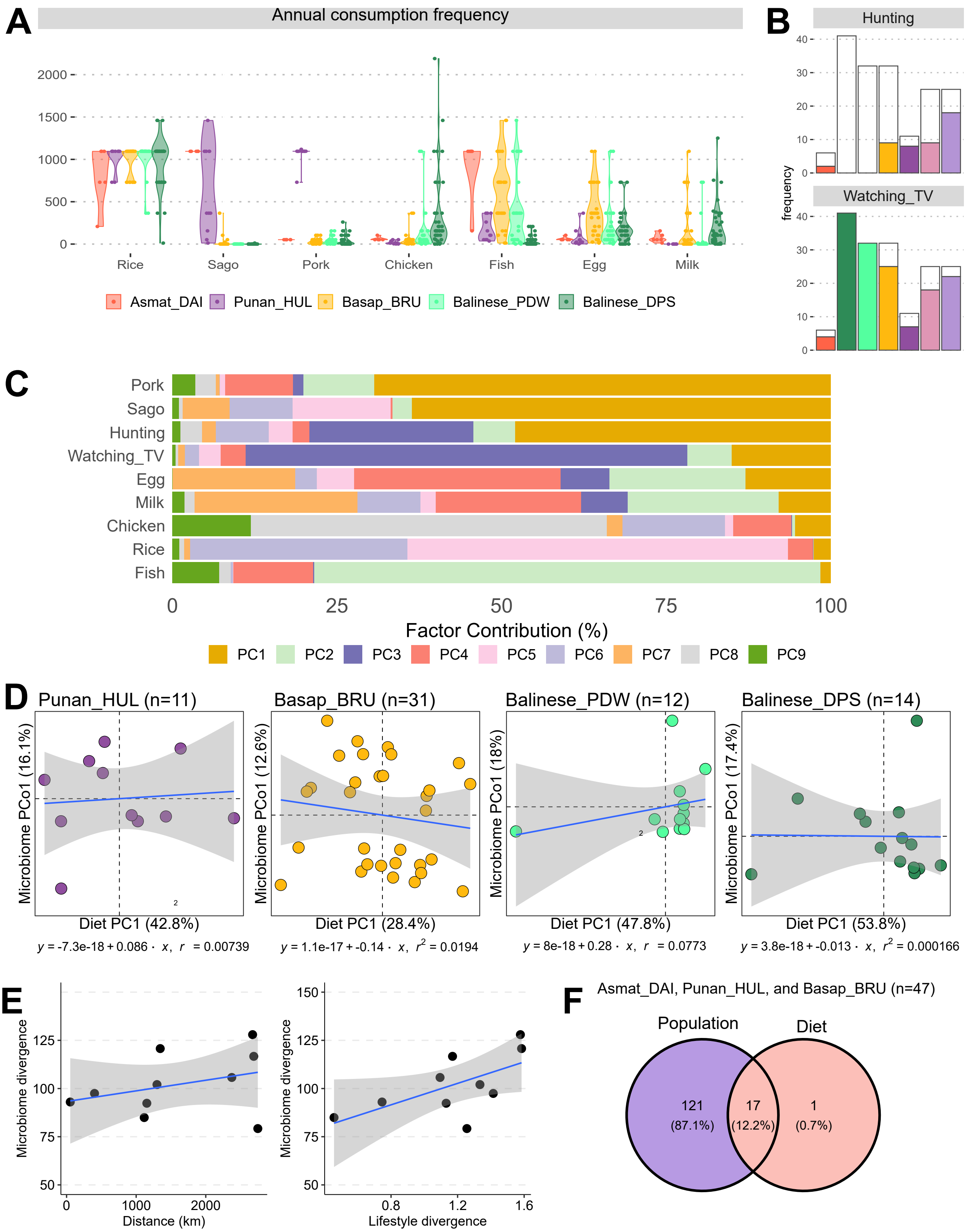


**Supplementary Figure S3. Lifestyle and dietary patterns across Indonesian samples.** **A)** Distribution of annual consumption frequencies for common dietary items by population, for 122 participants from five populations. **B)** Number of individuals from seven populations (from a total of 172 participants) who reported practising hunting and watching television at least once in the year leading to the time of enrolment. Bar outlines reflect population sample sizes. **C)** Loadings of the lifestyle PCA showing the proportional contributions of dietary components to principal components (PC1–PC9). This lifestyle PCA was performed on 73 Indonesian individuals with matching microbiome data (related to Figure 3). **D)** Per-population linear regression analyses of the first principal component (PC1) of the lifestyle PCA against the first principal coordinate (PCo1) of the microbiome PCoA (n = sample size). **E)** Microbiome divergence (pairwise Aitchison distance) versus geographic distance (great-circle distance, km; left) and lifestyle divergence ($\chi^2$ distance; right) for the five populations. **F)** Venn diagram comparing the number of MAGs significantly associated ($p < 0.05$) with diet and population across 3 populations (Asmat Daikot, Punan Respen Hulu, and Basap), as identified by MaAsLin2.

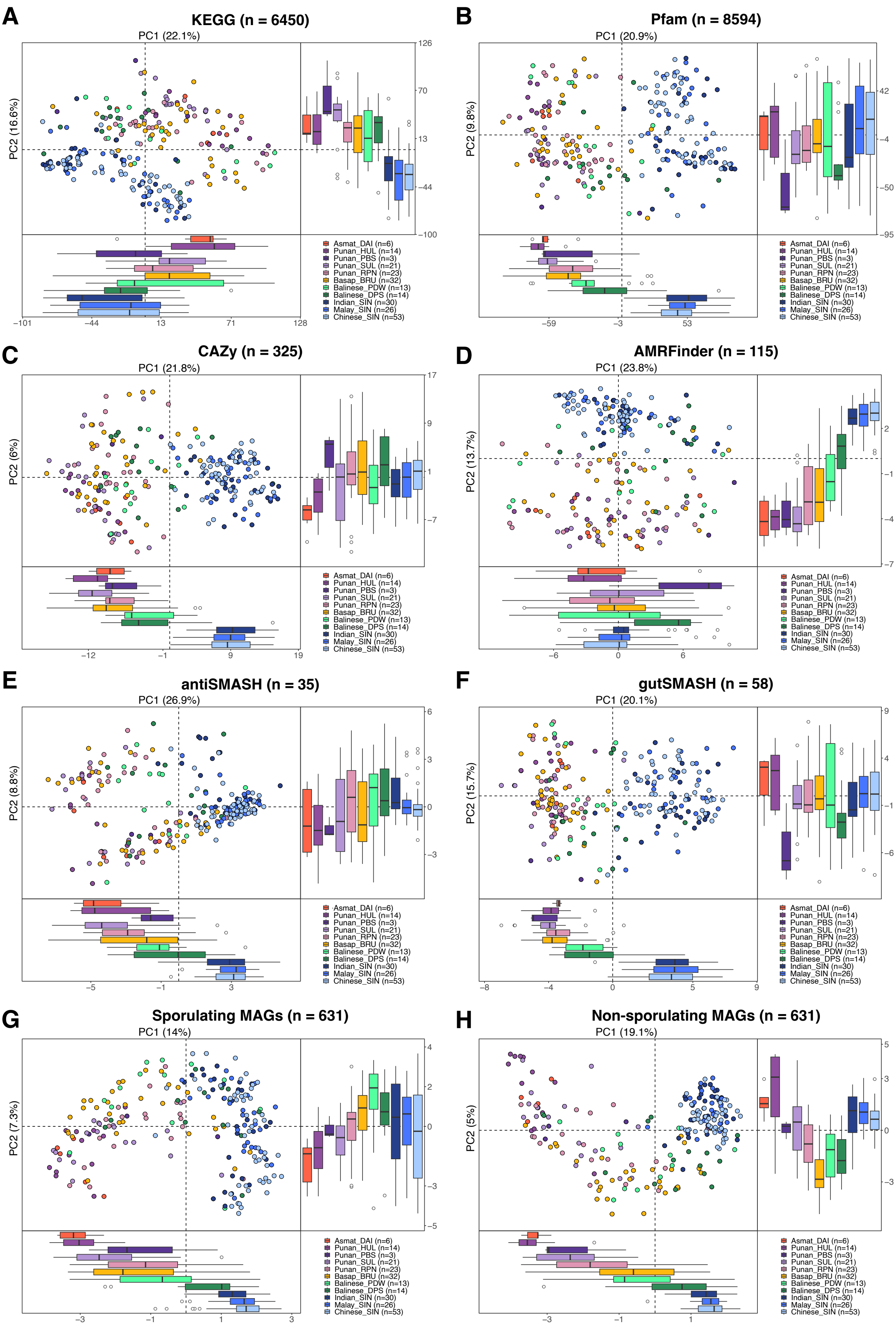


**Supplementary Figure S4. Microbiome functional and sporulation-associated structure.** Principal component analyses of **A**) KEGG functional orthologs, **B**) Pfam protein families, **C**) carbohydrate-active enzyme and substrates (CAZy), **D**) antimicrobial resistance genes (AMRFinder), **E**) secondary metabolite biosynthetic gene clusters (antiSMASH), **F**) primary metabolic gene pathways (gutSMASH), **G**) sporulating MAGs, and **H**) non-sporulating MAGs, of 116 Indonesian and 109 Singaporean individuals. Here, sporulating or non-sporulating is discriminated by the presence or absence of the master regulator gene essential for sporulation i.e., Spo0A. Colours denote population. Feature sample sizes are indicated in parentheses adjacent to headers and population sample sizes are indicated in parentheses in panel legends. Box and whiskers plots show the positional distribution of individuals by population along the first and second principal components. Centre lines denote medians, box edges indicate lower and upper quartiles, and whiskers extend to 1.5 times the interquartile range.

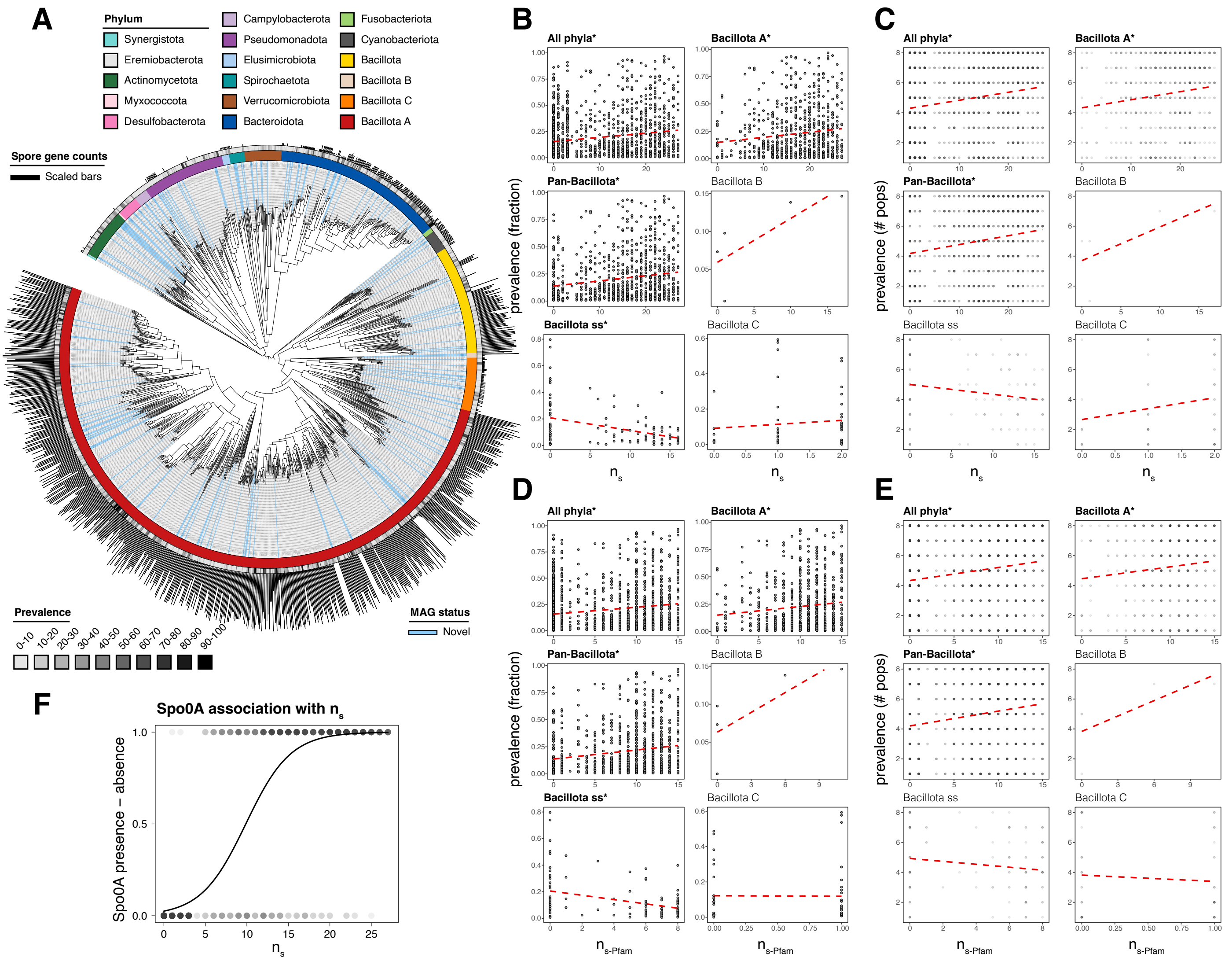


**Supplementary Figure S5. Correlation of sporulation gene counts with microbe prevalence.** **A**) Phylogenetic tree of IndoMEE MAGs annotated with phylum (inner coloured ring), novelty (blue radii), prevalence (outer grayscale ring), and number of KEGG-annotated sporulation genes $n_s$ (outer bars). **B**-**E**) Correlation of MAG sporulation gene count ($n_s$, $n_{s\text{-Pfam}}$) with prevalence, specifically overall prevalence fraction (**B**, **D**) and the number of populations in which the MAG is present (**C**, **E**); under KEGG (**B**, **C**) and Pfam (**D**, **E**) annotations. Significant correlations (i.e., FDR-adjusted $p < 0.05$) are indicated by bold letters and asterisk in panel headers. In **C** and **E**, circles with darker shades of grey reflect higher frequency of points. Best-fit regression lines are shown in red. **F**) Logistic regression of MAG sporulation gene counts ($n_s$) against the presence-absence of the Stage 0 sporulation protein A (Spo0A) across Bacillota phyla ($p < 0.001$, Nagelkerke's pseudo-$R^2 = 0.805$). Circles with darker shades of grey reflect higher frequency of points.

**Supplementary Table S1. Baseline community characteristics and lifestyle**

| Population (Code) | Sample size (n) | Age[a] (years) | BMI[b] | Sex M:F | Lifestyle Category | Location Name | Environment Type | Food Source | Latitude, Longitude |
|---|---|---|---|---|---|---|---|---|---|
| Asmat Daikot (Asmat_DAI) | 5 | 35 (28 - 51) | 20.3 (16.2 - 20.9) | 5:0 | remote | Daikot | riverside village | subsistence crops, fishing, market goods | -5.5086, 139.2380 |
| Punan Batu (Punan_PBS) | 3 | NA | NA | 2:1 | remote | Sajau | forest camps | gathered foods (fruit, vegetables, honey), frequent hunting, limited horticulture, market goods | 2.6258, 117.4019 |
| Punan Tubu Hulu (Punan_HUL) | 12 | 31 (22 - 55) | 22.2 (16.5 - 27.2) | 6:6 | remote | Rian Tubu | forest village | daily fishing, frequent hunting, subsistence crops, market goods | 3.3869, 116.0648 |
| Punan Aput (Punan_SUL) | 18 | 52 (35 - 70) | 23.05 (19.2 - 29.1) | 17:1 | remote | Long Sule | forest village | subsistence crops, fishing, occasional hunting | 1.80607, 115.5670 |
| Punan Tubu Respen (Punan_RPN) | 20 | 57.5 (25 - 78) | 20.8 (17 - 29.7) | 18:0 | rural | Respen Tubu | peri-urban village | subsistence crops, subsistence husbandry, market goods, occasional hunting | 3.7367, 116.4017 |
| Basap (Basap_BRU) | 31 | 36 (21 - 88) | 22.7 (17.9 - 31) | 9:22 | rural | Teluk Sumbang | coastal village | subsistence crops, fishing, market goods, minimal hunting | 1.0358, 118.8867 |
| Balinese Pedawa (Balinese_PDW) | 13 | 46.5 (27 - 53) | 27.4 (20.1 - 43.6) | 6:6 | rural | Pedawa | highland village | market goods, subsistence husbandry | -8.233010, 115.040000 |
| Balinese Denpasar (Balinese_DPS) | 14 | 21 (20 - 27) | 21.45 (18.9 - 54.1) | 0:14 | urban | Denpasar | coastal city | market goods | -8.6704, 115.2126 |
| All Samples | 116 | 42 (20 - 88)[c] | 22.4 (16.2 - 54.1)[d] | 63:50[e] | | | | | |

**a-b**, values are displayed as median (min - max). **c**, 7 NAs for age (3 Punan_PBS, 1 Punan_HUL, 1 Balinese_PDW). **d**, 8 NAs for BMI (3 Punan_PBS, 1 Punan_HUL, 3 Punan_RPN, 1 Balinese_PDW). **e**, 3 NAs for Sex (2 Punan_RPN, 1 Balinese_PDW).
